# Self-inactivating AAV-CRISPR at different ages enables sustained amelioration of Huntington’s disease deficits in BAC226Q mice

**DOI:** 10.1101/2025.06.24.661435

**Authors:** Yuanyi Dai, Zuliayeti Abudujielili, Yunyi Ding, Wanping Huang, Jianhang Yin, Liqiong Ou, Jiazhi Hu, Sushuang Zheng, Chenjian Li

## Abstract

Huntington’s disease (HD) is a monogenic autosomal dominant neurodegenerative disorder caused by a CAG repeat expansion in the first exon of the *HTT* gene, yielding a gain-of-toxic-function mutant Huntingtin protein mHTT. CRISPR/Cas9 is a potentially powerful therapeutic tool for treating HD by eliminating mutant *HTT* (m*HTT*) gene. We developed a specific SaCas9 guide RNA to target human m*HTT*, and a self-inactivating gene editing system that abolishes SaCas9 after a short transient expression for high gene editing efficiency and maximal safety to prevent off-target effects. Both conventional and the new self-inactivating gene editing systems achieved successful elimination of m*HTT* gene, 60-90% mHTT protein and 90% of mHTT aggregation in BAC226Q HD mouse brains, which resulted in significant long-term rescue of neural pathology, motor deficits, weight loss and shortened lifespan. These beneficial effects were observed when gene editing was applied before, at and well after the on-set of pathological and behavioral abnormalities. These proof-of-concept data demonstrate that gene editing can be a highly effective therapeutic approach for HD and other inherited neurodegenerative diseases.

**One Sentence Summary:** Self-inactivating CRISPR for mutant *huntingtin* in HD mice achieved long-term rescue of neural pathology, motor deficits, weight loss and survival.

## INTRODUCTION

This year, nervous system disorders passed cardiovascular diseases and cancer, and became the most prevalent illness of humanity (*1*), with neurodegenerative disorders as the major unmet medical and social needs in a rapidly aging population worldwide. In age-dependent, progressive and fatal disorders such as Alzheimer’s Disease (AD), Parkinson’s Disease (PD), Amyotrophic Lateral Sclerosis (ALS) and HD, the pathogenesis is extremely complex from molecule, cell, neural circuit, brain regional degeneration to behavior and symptoms. This provided numerous potential targets that attracted tremendous efforts from research groups and pharmaceutical industry to develop intervention strategies. However, to date, effective treatments to prevent the onset and progression of these diseases are elusive. All of these diseases have a strong genetic component, varying from substantial percentages to monogenic with near 100% penetrance. Many of gene mutations have been identified as the direct causes for specific disorders (*2*), and some are shared between hereditary and sporadic cases. Since gene mutations in these diseases are the most upstream events, strategies to correct mutations, delete disease genes, or add back absent genes are being pursued actively with huge potential.

Among these neurodegenerative diseases, HD is monogenic with near full penetrance by m*HTT*. HD is autosomal-dominant and caused by a CAG repeat expansion (≥36 repeats) in the first exon of the *huntingtin* gene (*3, 4*), which encodes a toxic gain-of-function mHTT protein with an expanded polyglutamine (polyQ) tract in the N-terminal region. mHTT can cause a wide range of cellular dysfunctions, which finally lead to specific neuronal loss, mainly in medium spiny neurons (MSNs) in striatum and pyramidal neurons in cortex (*5*). Since mHTT is the cause of HD and the severity of HD symptoms is linked to mHTT expression (*6, 7*), reducing mHTT is a valid and direct therapeutic strategy for HD patients.

HTT-lowering have been explored by multiple methods, including increasing protein degradation by enhancing proteasome or autophagy functions (*8*), as well as degrading RNA via antisense oligonucleotides (ASOs) (*9–11*), RNA interference (RNAi) (*12–18*) and CRISPR/Cas13 (*19, 20*). These strategies have been evaluated for safety, tolerability and efficacy in preclinical studies, and are being actively explored in clinical trials (*21*) (www.clinicaltrials.gov, identifier NCT02519036, NCT04120493). Furthermore, allele-selectively targeting *HTT* CAG repeats or single nucleotide polymorphism (SNP) has been attempted by Zinc-finger Nucleases (ZFNs) approach (*22–24*), and CRISPR-based approach (*20, 25–28*).

CRISPR/Cas9, conventional (*29–32*) or self-inactivating (*33–37*) forms, are powerful tools for correcting or deleting disease-causing mutated genes. Specifically for HD, the most upstream mHTT lowering could be achieved by CRISPR/Cas9-mediated DNA-targeting. Therapies targeting *HTT* DNA are expected to ameliorate all aspects of HD. Recent studies have shown the feasibility of CRISPR/Cas9-mediated *HTT* gene disruption in cell models (*25*), mouse models (*26, 27, 34, 38, 39*) and large mammals (*40*). Several of above studies (*9, 12, 18, 38, 39, 41*), along with previous findings (*42–45*) suggested that significant suppression of wild-type *HTT* in adult mouse, non-human primates (NHP) and human might be tolerated. To date, none of these CRISPR/Cas9-based studies entered clinical trials yet.

To better push forward the CRISPR/Cas9-mediated gene editing into clinical trials, we developed a self-inactivating CRISPR/Cas9 system to limit Cas9 expression only transiently for maximal safety in HD therapeutics. To verify the feasibility of CRISPR/Cas9-mediated gene therapy for HD, we tested in a robust and suitable mouse model BAC226Q that expresses full-length human m*HTT* with a stable 226 mixed CAG-CAA repeats beyond the 150 CAGs threshold for toxicity (*46, 47*). BAC226Q recapitulates main aspects of HD pathology and symptoms, including age-dependent and progressive neural pathology, cytosolic and nuclear mHTT aggregation in striatal and cortical neurons, motor deficits, weight loss and shortened lifespan (*48*).

Both conventional and self-inactivating CRISPR/Cas9 efficiently and permanently disrupted the human m*HTT* transgene, reduced 60-90% of mutant HTT protein and 90% of cytosolic aggregates and nuclear inclusions, which led to significant rescue of the above mentioned deficits in BAC226Q mice. We also tested the effectiveness of gene editing at various mouse ages corresponding to human patient pre-symptomatic, onset and symptomatic disease stages.

## RESULTS

### CRISPR/Cas9 design and validation of the cleavage of human *HTT* in cells

For complete human m*HTT* elimination, sgRNAs-HTTg1 and HTTg2 were designed to target the human *HTT* N-terminal 17-amino-acid domain (N17)-coding DNA sequence before CAG repeats (**Fig. 1A**). N17 is a conserved region across most of the species and is shown to play an important role in mHTT-induced pathogenesis (*49*). For *in vivo* experiments, cytomegalovirus (CMV)-driven SaCas9 and U6-driven *HTT* sgRNA were inserted into a single AAV vector (**Fig. 1B**). In HEK293T cells expressing human HTT-exon 1 fragment with 20Q or 120Q and in-frame N-terminal enhanced green fluorescent protein fusion (EGFP-hHTT-20Q or EGFP-hHTT-120Q), both sgRNAs resulted in robust HTT reduction, with the better performing SaCas9-HTTg1 decreasing HTT by 70%-80%, and therefore chosen for future experiments (**Fig. 1, C to K**).

**Fig. 1.**
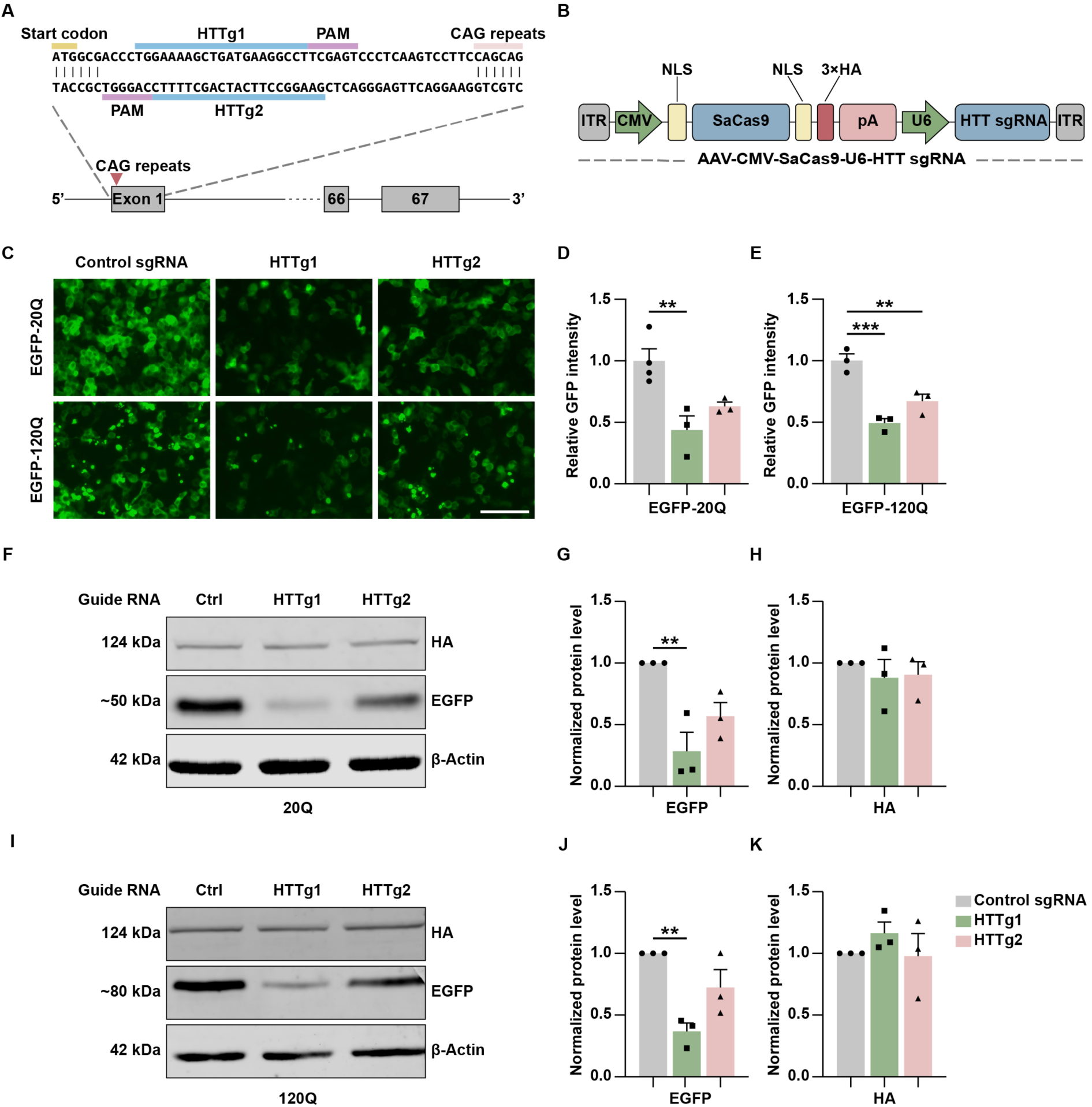
Design and cleavage efficiency of human *HTT* of AAV-CRISPR/Cas9. (**A**) Schematic representation of targeted human HTT exon 1. sgRNAs HTTg1 and HTTg2 were designed to target the N17 domain-coding DNA sequence, which is located upstream of the CAG repeats. PAM, protospacer-adjacent motif. (**B**) Schematic representation of AAV-SaCas9-HTT sgRNA vector. U6-driven sgRNA and a CMV-driven Cas9 from *Staphylococcus aureus* tagged with HA were packaged into a single AAV vector of 4.5 kb. ITR, inverted terminal repeat; NLS, nuclear localization signal sequence; 3×HA, three tandem repeats of the human influenza hemagglutinin (HA) tag. (**C** to **K**) Validation of human HTT editing in HEK293T cells expressing EGFP-hHTT-20Q or EGFP-hHTT-120Q reporter system. (C) SaCas9-HTTg1 and SaCas9-HTTg2 reduced EGFP-fusion protein expression and aggregation in cells, whereas the control SaCas9-sgRNA did not. Scale bar: 100 μm. (D and E) Quantitative analyses of EGFP-20Q (D) or EGFP-120Q (E) fluorescence intensity. \*\**p*=0.0083 (20Q), \*\**p*=0.0087 (120Q), \*\*\**p*=0.0009 (120Q). (F to K) Western-blot detecting EGFP and SaCas9 (anti-HA) protein level in EGFP-hHTT-20Q (F to H) or EGFP-hHTT-120Q (I to K) reporter system. (F and I) SaCas9-HTTg1 and SaCas9-HTTg2 reduced EGFP-hHTT protein expression compared to control SaCas9-sgRNA with equivalent SaCas9 expression level. (G, H, J and K) Quantification of relative EGFP (G and J) and SaCas9 (H and K) protein expression. N=3 per group; \*\**p*=0.0087 (20Q), \*\**p*=0.0069 (120Q). Data were shown as mean ± SEM. One-way ANOVA with Tukey’s *post hoc* multiple comparisons test.

### SaCas9-HTTg1-mediated human mHTT disruption in BAC226Q mice

To evaluate the effects of CRISPR/Cas9-mediated m*HTT* gene editing *in vivo*, we used BAC226Q expressing full-length human m*HTT* with 226 CAG-CAA repeats (*48*). This mouse model recapitulates a full-spectrum of age-dependent and progressive HD-like phenotypes without unwanted and erroneous phenotypes, and therefore a powerful tool for testing gene-modifying therapeutics.

A longitudinal study of biochemistry, cellular pathology and behavioral tests was designed in BAC226Q mice with AAV-SaCas9-HTTg1 (**Fig. 2A**). To evaluate AAV delivery system, we first confirmed that AAV9 stereotactic injection into striatum and motor cortex, the two major HD pathology areas, yielded highly efficient neuronal transduction without inducing immunoreactivity (**fig. S1, A to H**). Subsequently, AAV-SaCas9-HTTg1 was injected into mouse brains at the age of 1 month, resulting in successful deletion of m*HTT* gene and life-long 60-90% reduction of mHTT protein (**Fig. 2, B and C**). In injected areas of striatum and cortex, AAV-SaCas9-HTTg1 reduced mHTT near completion (**Fig. 2D**). Furthermore, among cells effectively transduced by AAV-SaCas9-HTTg1, 94-96% mHTT was eliminated (**fig. S2, A to D**). We further investigated the consequence of effective gene editing on mHTT neuropil aggregates and nuclear inclusions, cardinal HD pathological hallmarks (*50*), which were accurately recapitulated in BAC226Q mice (*48*). By 4 months of age, BAC226Q mice exhibit clear cytoplasmic and nuclear mHTT aggregation in striatum and mainly cytoplasmic aggregation in cortex. As pathology progresses to 7 months, mHTT nuclear inclusions become more prominent (**Fig. 2, E to L and fig. S3, A and B**). In 4-month-old BAC226Q mice, AAV-SaCas9-HTTg1 reduced total mHTT aggregation signal in striatal and cortical neurons by 90% and 55% respectively (**Fig. 2, E to H**). At 7 months, the rescue effects remained robust, with total mHTT reduction of 86% in striatum and 48% in cortex (**Fig. 2, I to L**). We further investigated the reduction of cytoplasmic and nuclear mHTT aggregates in number and size separately, and observed the same robust reduction (**fig. S3, A to J**). The percentage of nucleus with mHTT aggregates were both reduced in striatum and cortex (**fig. S3, K to L**). Interestingly, AAV-SaCas9-HTTg1 did not change GFAP-immunoreactivity in BAC226Q mice (**fig. S4, A and B**). In conclusion, these results confirmed a sustained and remarkable reduction of mHTT protein and aggregates *in vivo* for lifetime, just by a single dose of AAV-SaCas9-HTTg1.

**Fig. 2.**
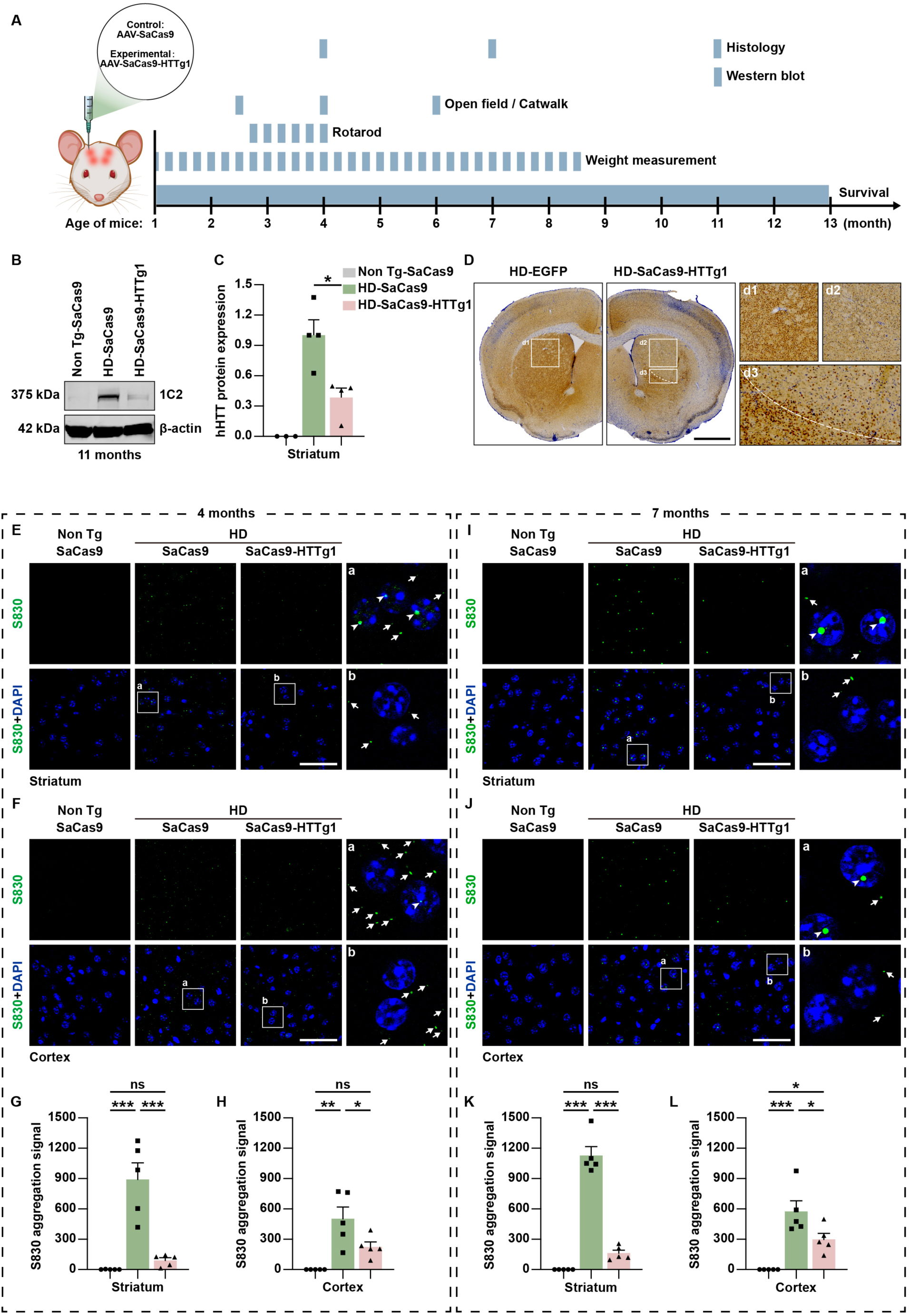
Significant reduction of human mHTT by intrastriatal and intracortical AAV9-mediated SaCas9-HTTg1 editing in BAC226Q mice. (**A**) Experimental timeline and corresponding analyses for *in vivo* study. (**B** to **D**) Significant reduction of mHTT protein in the striatum and cortex of BAC226Q mouse injected with AAV-SaCas9-HTTg1. Non Tg-SaCas9, non-transgenic mice injected with AAV-SaCas9; HD-SaCas9, BAC226Q mice injected with AAV-SaCas9; HD-SaCas9-HTTg1, BAC226Q mice injected with AAV-SaCas9-HTTg1. (B and C) Western-blot analysis (B) and quantification (C) of sustained decrease of striatal mHTT protein in HD-SaCas9-HTTg1 mouse brain at 11 months. 1C2 antibody was used to detect mHTT protein with β-actin as a loading control. N=3 mice for Non Tg-SaCas9, N=4 for HD-SaCas9 and HD-SaCas9-HTTg1; \**p*=0.0104. (D) Immunohistochemistry of significant mHTT decrease in striatal virus-diffused areas in 11-month-old HD-SaCas9-HTTg1 mouse. S830 antibody was used to detect mHTT. AAV9-EGFP was a control. AAV9 diffusion boundary was indicated by dotted lines. Enlarged figures were shown on the right. Scale bar: 2000 μm. (**E** to **L**) Significant decrease of S830-positive mHTT aggregates in HD-SaCas9-HTTg1 mice. (E, F, I and J) Immunofluorescent histology of mouse striatum (E and I) and primary motor cortex (F and J) at 4 months (E and F) and 7 months (I and J), boxed areas were enlarged on the right. Cytoplasmic aggregates (arrows) and nuclear inclusion (arrowheads). mHTT (green) and nucleus (blue) were detected by S830 and DAPI, respectively. Scale bar: 50 μm. (G, H, K and L) Quantification of mHTT aggregation signal in striatum (G and K) and primary motor cortex (H and L) at 4 months (G and H) and 7 months (K and L). Aggregation signal was analyzed by particle analysis in ImageJ and calculated by the number of aggregates × integrated density. N=5 mice per group; For 4 months, *** *p*<0.0001 (Non Tg-SaCas9 versus HD-SaCas9, striatum), \*\*\**p*=0.0002 (HD-SaCas9 versus HD-SaCas9-HTTg1, striatum), \*\**p*=0.0011, \**p*=0.0495, ns=0.7886 (striatum), ns=0.1176 (cortex); For 7 months, \*\*\**p*<0.0001 (striatum), ns=0.1211 (striatum), \*\*\**p*=0.0002 (cortex), \**p*=0.0265 (Non Tg-SaCas9 versus HD-SaCas9-HTTg1, cortex), \**p*=0.0402 (HD-SaCas9 versus HD-SaCas9-HTTg1, cortex). Data were shown as mean ± SEM. One-way ANOVA with Tukey’s *post hoc* multiple comparisons test.

### Robust and long-lasting rescue of motor deficits, weight loss and lifespan by CRISPR/Cas9-mediated m*HTT* gene deletion

Since AAV-SaCas9-HTTg1 effectively deleted m*HTT* gene, decreased mHTT protein and aggregates, we set out to investigate whether gene editing rescues cardinal HD-like phenotypes such as age-dependent motor deficits, weight loss and shortened lifespan in BAC226Q mice (*48*).

BAC226Q mice exhibit normal motor function until 2-3 months of age, followed by severe hyperactivity and chorea-like movement from 3-4 months, and a subsequent reduction in motor activity from 6-7 months. These deficits and rescue effects of AAV-SaCas9-HTTg1 were observed in home cage behavior (**movie S1**) and analyzed by catwalk, rotarod and open-field tasks. In catwalk task, BAC226Q mice developed progressive gait irregularities reflected by scrambled step sequences and footprint area reductions (**Fig. 3, A to E**). Severe scrambled step sequences at 6 months were fully rescued by AAV-SaCas9-HTTg1 (**Fig. 3, A and B and movie S2**). Decreased footprint areas at 2.5 months, 4 months, and 6 months in BAC226Q mice were significantly ameliorated by AAV-SaCas9-HTTg1 (**Fig. 3, C to E**). In rotarod task, BAC226Q mice exhibited strong deficits in motor coordination and balance measured by latency to fall, which were significantly improved in HD-SaCas9-HTTg1 mice (**Fig. 3F**). In open-field test, non-transgenic control mice travelled across the cage, whereas BAC226Q mice circled in periphery. This deficit was reversed in HD-SaCas9-HTTg1 mice (**Fig. 3G**). Furthermore, dramatically increased total travel distance in BAC226Q mice was attenuated by about 30-35% (**Fig. 3H**).

**Fig. 3.**
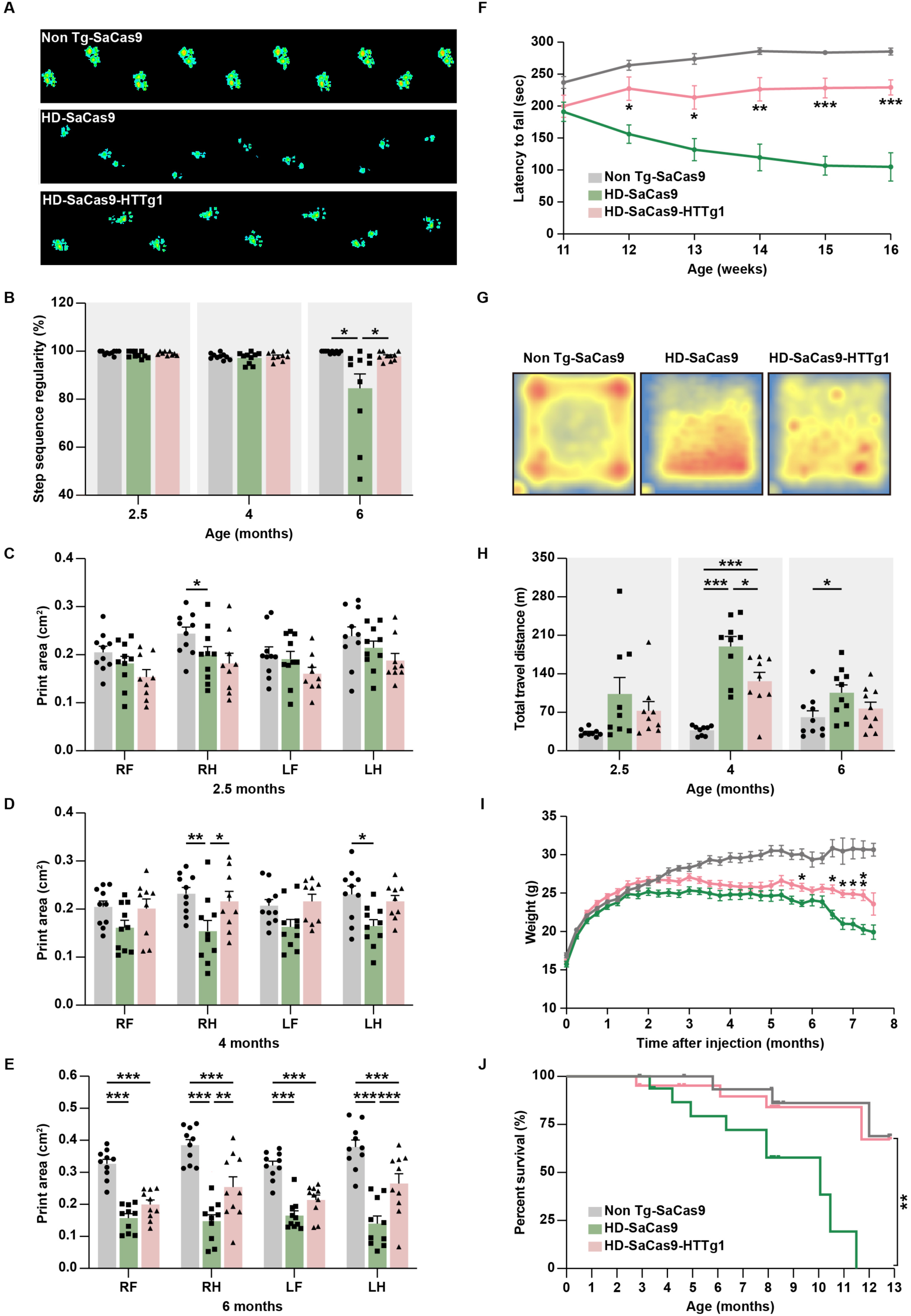
Robust and long-lasting motor performance improvement, weight loss attenuation and lifespan rescue by gene editing in BAC226Q mice. (**A** to **E**) Improved gait performance in HD-SaCas9-HTTg1 mice analyzed by catwalk. (A) Representative footprints of Non tg-SaCas9 (upper), HD-SaCas9 (middle) and HD-SaCas9-HTTg1 mice (lower) at 6 months. (B) Significant increase of step sequence regularity index in 6-month HD-SaCas9-HTTg1 mice. N=10 mice per group; \**p*=0.0309 (HD-SaCas9 versus HD-SaCas9-HTTg1); One-way ANOVA with Tukey’s *post hoc* multiple comparisons test. (C to E) Footprint area quantification of Non tg-SaCas9, HD-SaCas9 and HD-SaCas9-HTTg1. (C) Similar baselines at 2.5 months. (D and E) Deficits in HD-SaCas9 were significantly rescued in HD-SaCas9-HTTg1 at 4 months (\**p*=0.0231, RH) and 6 months (\*\**p*=0.0010, RH; \*\*\**p*<0.001, LH). N=9∼10 mice per group. Two-way ANOVA with Tukey’s *post hoc* multiple comparisons test. RF, right front; RH, right hind; LF, left front; LH, left hind. (**F**) Improved coordination and balance ability in rotarod test in HD-SaCas9-HTTg1 mice. N=9∼10 mice per group; HD-SaCas9-HTTg1 versus HD-SaCas9, \**p*=0.0202 (12 weeks), \*\**p*=0.0126 (13 weeks), \*\**p*=0.0035 (14 weeks), \*\*\**p*<0.001 (15 and 16 weeks); Two-way ANOVA with Tukey’s *post hoc* multiple comparisons test. (**G** and **H**) Rescue of locomotor performance in HD-SaCas9-HTTg1 mice in open-field test. (G) Representative heatmaps exhibiting time and location to stay for 6-month-old Non Tg-SaCas9 (left), HD-SaCas9 (middle) and HD-SaCas9-HTTg1 (right) mice. Red color represents cumulative appearance. (H) Total travel distance of 2.5, 4 and 6-month mice. N=9∼10 mice per group; HD-SaCas9-HTTg1 versus HD-SaCas9, \**p*=0.0109 (4 months), \**p*=0.0456 (6 months); One-way ANOVA with Tukey’s *post hoc* multiple comparisons test. (**I**) Weight loss attenuation in HD-SaCas9-HTTg1 mice. Non Tg-SaCas9 N=27; HD-SaCas9 N=26; BAC226Q-HTTg1 N=26; \**p*<0.05, \*\**p*<0.01; Two-way ANOVA with Tukey’s *post hoc* multiple comparisons test. (**J**) Complete rescue of survival time by gene editing. The survival of HD-SaCas9-HTTg1 mice was significantly improved compared with HD-SaCas9 and reached to a level consistent with that of Non Tg-SaCas9. *p*=0.0061 (HD-SaCas9-HTTg1 versus HD-SaCas9), *p*=0.7434 (HD-SaCas9-HTTg1 versus Non Tg-SaCas9), *p*=0.0025 (HD-SaCas9 versus Non Tg-SaCas9). Non Tg-SaCas9 (N=17), HD-SaCas9 (N=17) and HD-SaCas9-HTTg1 (N=21) mice. Log-rank (Mantel-Cox) test. Data were shown as mean ± SEM.

BAC226Q mouse model recapitulates HD-like age-dependent progressive weight loss and shortened lifespan as in HD patients (*48*). AAV-SaCas9-HTTg1 attenuated body weight loss by 5-22% (**Fig. 3I**), and promoted lifespan dramatically (**Fig. 3J**).

Taken together, significant and long-term rescue of motor deficits, weight loss and shortened lifespan were obtained if gene editing eliminated m*HTT* gene in 1-month-old BAC226Q mice before any pathology and motor deficits were observed.

### Rescue effects of gene editing at different pathogenic time points

An important question relevant to clinical treatment is at which disease stages gene editing is still effective (*51*). BAC226Q mice accurately recapitulated the age-dependent progression from normal development, to phenotype onset and worsening, and severe end stages. Therefore, we investigated the rescue effects of gene editing in BAC226Q at 1, 4 and 7 months of age, which are before, at and well after the on-set of pathological and behavioral abnormalities (**Fig. 4A**).

**Fig. 4.**
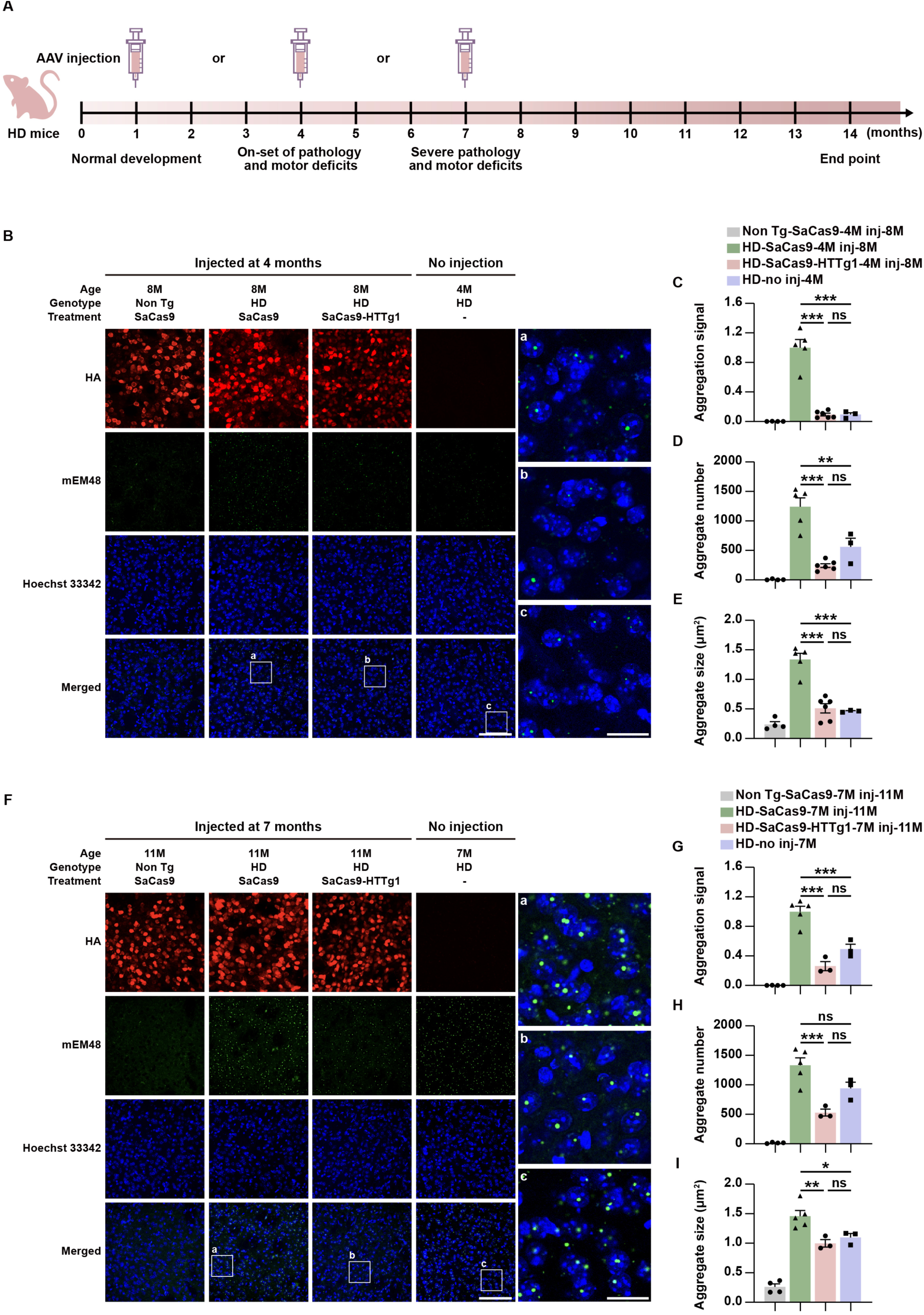
Robust reduction of human mHTT aggregation by gene editing at 4 and 7 months. (**A**) Experimental timeline in HD mice and corresponding phenotypes at different time points. (**B** to **I**), Significant reduction of mEM48-positive mHTT aggregates (green) in the striatum from 4-month-injected HD mice (B to E) and 7-month-injected HD mice (F to I). Mouse brain samples were collected 4 months post injection. (B and F) Representative figures showing that mHTT aggregation was decreased in 4-month-injected HD-SaCas9-HTTg1 mice (B) and 7-month-injected HD-SaCas9-HTTg1 mice (F). mHTT (green) and SaCas9 (red) were detected by mEM48 and anti-HA antibody, respectively. Scale bar: 100 μm. Enlarged figures were shown on the right side. Scale bar: 20 μm. (C to E and G to I) Quantification of mHTT aggregation signal (C and G), aggregate number (D and H) and aggregate size (E and I) in the striatum in 4-month-injected HD-SaCas9-HTTg1 mice (C to E) and 7-month-injected HD-SaCas9-HTTg1 mice (G to I). \*\*\**p*<0.0001 (C to E), \*\**p*=0.0031 (D), \*\*\**p*=0.0006 (G, HD-no inj-7M versus HD-SaCas9-7M inj-11M), \*\*\**p*<0.0001 (G, HD-SaCas9-7M inj-11M versus HD-SaCas9-HTTg1-7M inj-11M), \*\*\**p*=0.0007 (H), \*\**p*=0.0082 (I) and \**p*=0.0363 (I). Data were shown as mean ± SEM. One-way ANOVA with Tukey’s *post hoc* multiple comparisons test.

Gene editing at the first time point of 1 month in pre-phenotypic mice yielded strong benefits as shown above (**Fig. 2**, **Fig. 3, fig. S2 and Fig. 3**).

The second time point of gene editing was at 4 months when BAC226Q mice start to exhibit early HD-like phenotypes including significant mHTT aggregation and motor behavior abnormalities. We injected AAV-SaCas9-HTTg1 at 4 months and examined the rescue effects of brain pathology and behavior throughout the lifespan of mice. At 8 months, AAV-SaCas9-HTTg1 reduced overall mHTT aggregation signal by 91% (**Fig. 4, B and C**). Further detailed analyses revealed that the reduction of aggregate number and size were 80% and 62% respectively (**Fig. 4, D and E**). Importantly, the dramatic 10-fold increase of aggregations from 4 months to 8 months in BAC226Q was completely abolished by AAV-SaCas9-HTTg1 (**Fig. 4, B to E**). The similar mHTT aggregation reduction was maintained at 11 months (**fig. S5, A and B**). In motor behavioral tests, gait irregularities at 6 months (**fig. S6 A and B and movie S3**) and locomotor activity at 8 months (**fig. S6, C and D**) were both rescued, although slightly less effectively than that by 1-month injection.

The third time point of gene editing was at 7 months when BAC226Q mice show severe motor deficits and pathology such as bigger and more predominant mHTT nuclear inclusions. This intervention still robustly reduced overall mHTT aggregation signal by 74%, aggregation number by 60% and aggregation size by 32% (**Fig. 4, F to I**). Similar to the 4-month injection data, AAV-SaCas9-HTTg1 completely abolished the 1-fold increase of aggregations from 7 months to 11 months in BAC226Q (**Fig. 4, F to I and fig. S5, C and D**). Consistent with the robust rescue in pathology, BAC226Q post-injection survival time was greatly increased (**fig. S6E**). In open-field test, locomotor abnormality was not rescued (**fig. S6F**). Gait irregularities measured by catwalk test could not be performed due to severe motor deficits.

It is important that these data collectively demonstrated that gene editing at different pathogenic time points resulted in significant mHTT reduction and recovery of lifespan in all cases, and rescue of motor deficits in early injections.

### Highly effective and sequential deletion of mHTT and Cas9 by self-inactivating CRISPR/Cas9 rescued deficits in motor function and lifespan

To address the clinically relevant and general concern that conventional long-term Cas9 expression in patients, potentially for decades, may cause cumulative off-target effects and acquired immune responses against Cas9 harboring cells, it will be greatly advantageous to develop a strategy that eliminates both the target gene and subsequently Cas9 itself. Here, we report such a self-inactivating system that carries AAV-SaCas9-HTTg1 and an additional AAV-sgRNA to target Cas9 (gCas9), so that Cas9 sequence will be destroyed after *HTT* disruption (**Fig. 5, A and B**).

**Fig. 5.**
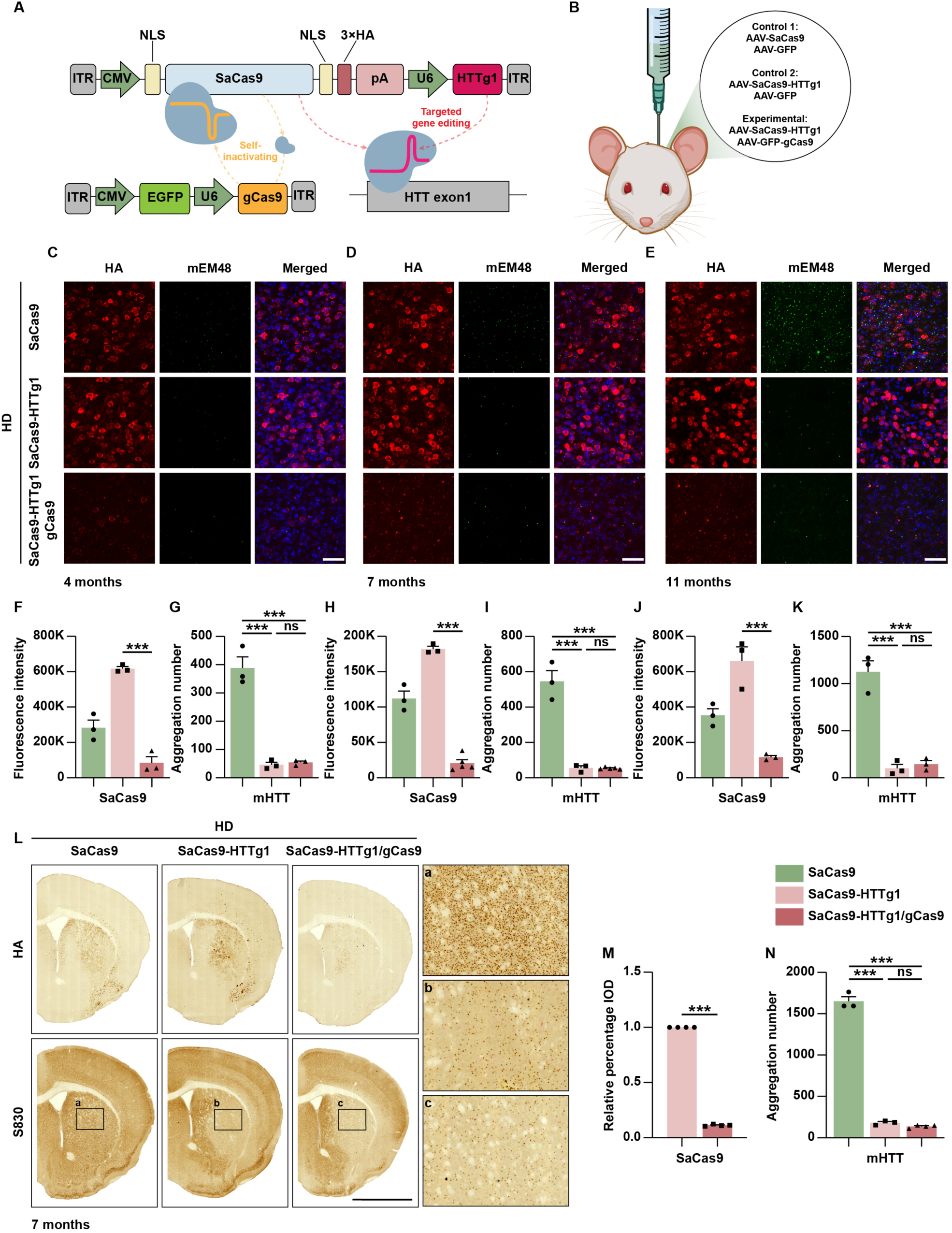
Self-inactivating gene editing system with high efficiency in reducing human mHTT and Cas9 *in vivo*. (**A**) Schematic representation of dual vectors for self-inactivating system. In AAV-SaCas9-HTTg1 vector, U6 drives HTTg1 to target human *HTT* and CMV drives SaCas9 gene. In AAV-EGFP-gCas9 vector, U6 drives gCas9 to target SaCas9 gene and CMV drives *EGFP* as a reporter gene. (**B**) Schematic diagram of dual AAV vectors delivered by intraparenchymal injection into striatum and primary motor cortex of 1-month-old BAC226Q and control littermates. (**C** to **N**) Self-inactivating system AAV-SaCas9-HTTg1/gCas9 robustly decreased SaCas9, and reduced mHTT aggregation with the same efficiency as the conventional system AAV-SaCas9-HTTg1 in BAC226Q mice. Representative immunofluorescence images of mEM48-labeled mHTT aggregation and HA-labeled SaCas9 in striatum of BAC226Q mice at 4 months (C), 7 months (D) and 11months (E). Scale bar: 50 μm. (F and G) Quantification of SaCas9 fluorescence intensity (F) and mHTT aggregation number (G) in (C). For SaCas9: N=3 mice per group; \*\*\**p*<0.0001. For mHTT: N=3 mice per group; \*\*\**p*<0.0001 (HD-SaCas9 versus HD-SaCas9-HTTg1), \*\*\**p*<0.0001 (HD-SaCas9 versus HD-SaCas9-HTTg1/gCas9), ns=0.9679. (H and I) Quantification of SaCas9 fluorescence intensity (H) and mHTT aggregation number (I) in (D). For SaCas9: N=3/5 mice per group; \*\*\**p*<0.0001. For mHTT: N=3/5 mice per group; \*\*\**p*<0.0001 (HD-SaCas9 versus HD-SaCas9-HTTg1), \*\*\**p*<0.0001 (HD-SaCas9 versus HD-SaCas9-HTTg1/gCas9), ns=0.9987. (J and K) Quantification of SaCas9 fluorescence intensity (J) and mHTT aggregation number (K) in (E). For SaCas9: N=3 mice per group; \*\*\**p*=0.0007. For mHTT: N=3 mice per group; \*\*\**p*=0.0002 (HD-SaCas9 versus HD-SaCas9-HTTg1), \*\*\**p*<0.0002 (HD-SaCas9 versus HD-SaCas9-HTTg1/gCas9), ns=0.9096. (L) Immunohistochemistry of HA-labeled SaCas9 (upper) and S830-labeled mHTT (lower) in 7-month-old BAC226Q mice. Boxed areas were enlarged on the right respectively. Scale bar: 2000 μm. (M) Quantification of SaCas9 reduction in (L). N=4 mice per group; \*\*\**p*<0.0001. (N) Quantification of striatal mHTT aggregation number in boxed areas of (L). N=3/4 mice per group; \*\*\**p*<0.0001 (HD-SaCas9 versus HD-SaCas9-HTTg1), \*\*\**p*<0.0001 (HD-SaCas9 versus HD-SaCas9-HTTg1/gCas9), ns=0.5473. Data were shown as mean ± SEM. One-way ANOVA with Tukey’s *post hoc* multiple comparisons test and unpaired two-tailed *t*-test.

To select the most effective gCas9, we used HEK293T cells expressing EGFP-hHTT-20Q/120Q to test 10 gCas9 against different SaCas9 coding sequences (**fig. S7A and table S1**). The results showed that gCas9-1 exhibited the highest efficiency in eliminating Cas9 without impairing mHTT reduction (**fig. S7, B to L**), and thus was used for subsequent *in vivo* experiments.

We then tested in BAC226Q mice. Similar to the conventional gene editing, the self-inactivating AAV-SaCas9-HTTg1/gCas9 effectively reduced mHTT by 94% at 4 months, 92% at 7 months and 92% at 11 months (**Fig. 5, C to N and fig. S8, A to F**). Meanwhile, AAV-SaCas9-HTTg1/gCas9 reduced above 90% Cas9 by 1 week post injection (**fig. S8, G and H**) and lasted throughout lifespan (**Fig. 5, C to N and fig. S8, A to F**). In mice injected with AAV9-SaCas9-HTTg1/AAV9-EGFP-gCas9, 93% of GFP-positive cells expressed neither SaCas9 nor mHTT (**fig. S9, A and B**). These data demonstrated that the self-inactivating system can efficiently eliminate SaCas9 without affecting the clearance of mHTT, and that Cas9 mediated gene editing events in neurons happen very rapidly well within 1 week post injection, after which Cas9 is no longer needed.

Further analyses revealed that AAV-SaCas9-HTTg1/gCas9 and AAV-SaCas9-HTTg1 injected mice showed equivalent rescue of step sequence irregularity (**Fig. 6, A and B and movie S4**), footprint area reduction (**Fig. 6C**) in catwalk task, locomotor hyperactivity in open-field test (**Fig. 6D**), latency to fall in rotarod test (**Fig. 6E**), and shortened lifespan (**Fig. 6F**).

**Fig. 6.**
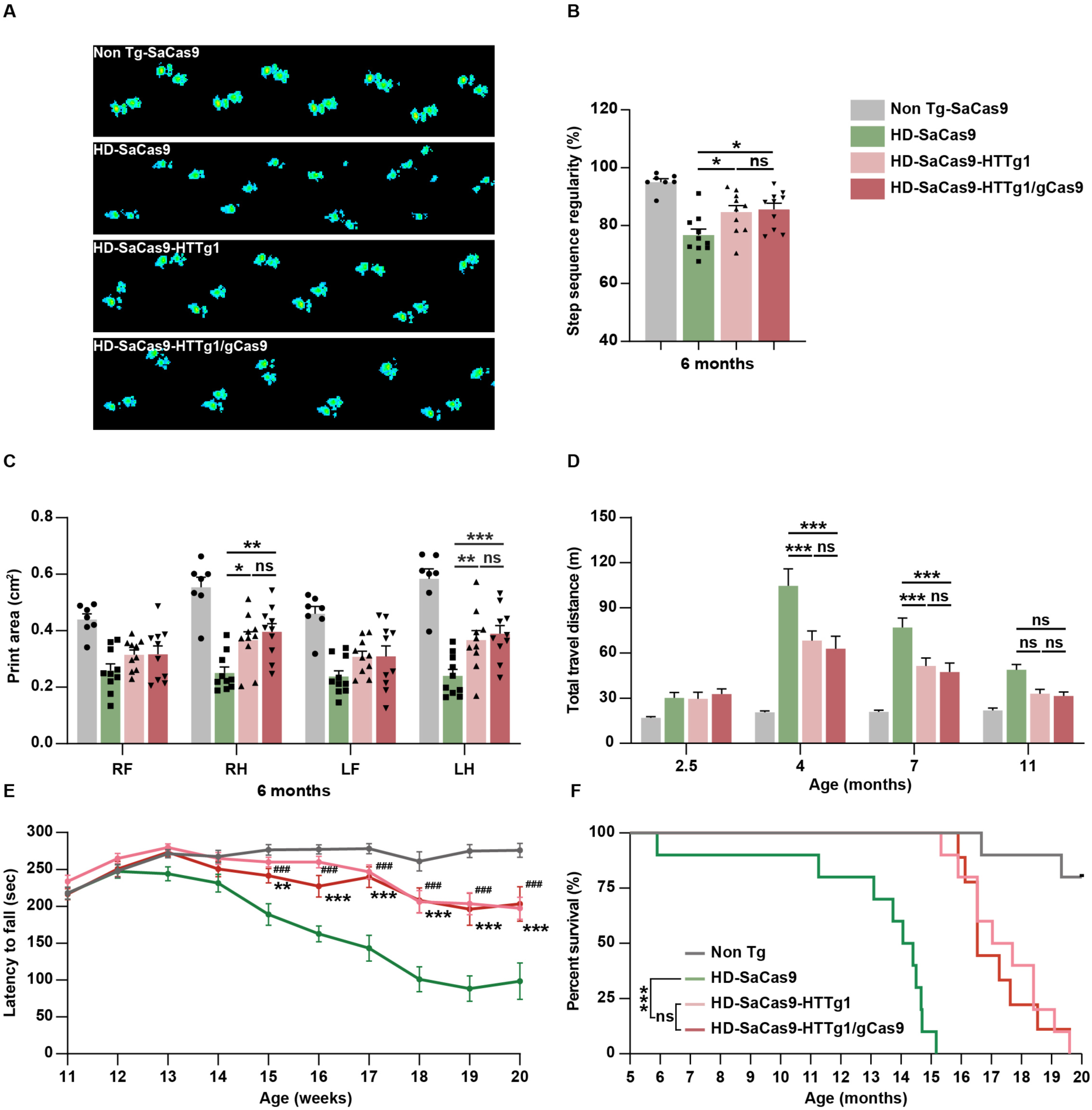
Successful improvement of motor performance and rescue of lifespan by self-inactivating system similar to conventional CRISPR/Cas9 in BAC226Q mice. (**A** to **C**) Self-inactivating AAV-SaCas9-HTTg1/gCas9 improved gait performance in catwalk analysis of BAC226Q mice at 6 months. (A) Representative footprints of Non tg-SaCas9, HD-SaCas9, HD-SaCas9-HTTg1 and HD injected with AAV-SaCas9-HTTg1/gCas9 (HD-SaCas9-HTTg1/gCas9) from top to bottom. (B) Increased step sequence regularity index in HD-SaCas9-HTTg1/gCas9 and HD-SaCas9-HTTg1 mice. N=10 mice per group; \**p*=0.0387 (HD-SaCas9 versus HD-SaCas9-HTTg1), \**p*=0.0164 (HD-SaCas9 versus HD-SaCas9-HTTg1/gCas9), ns=0.9844. One-way ANOVA with Tukey’s *post hoc* multiple comparisons test. (C) Quantification of footprint area in gait analysis. N=10 mice per group; \**p*=0.0136 (RH), \*\**p*=0.0010 (RH), ns=0.8595 (RH). \*\**p*=0.0057 (LH), \*\*\**p*=0.0007 (LH), ns=0.9319 (LH). Two-way ANOVA with Tukey’s *post hoc* multiple comparisons test. RF, right front; RH, right hind; LF, left front; LH, left hind. (**D**) Rescue of locomotor activity in HD-SaCas9-HTTg1/gCas9 and HD-SaCas9-HTTg1 mice in open-field test. N=15 mice per group; At 4 months: \*\*\**p*<0.0001 (HD-SaCas9 versus HD-SaCas9-HTTg1), \*\*\**p*<0.0001 (HD-SaCas9 versus HD-SaCas9-HTTg1/gCas9), ns=0.8905. At 7 months: \*\*\**p*=0.0003 (HD-SaCas9 versus HD-SaCas9-HTTg1), \*\*\**p*<0.0001 (HD-SaCas9 versus HD-SaCas9-HTTg1/gCas9), ns=0.9207. At 11 months: ns=0.1145 (HD-SaCas9 versus HD-SaCas9-HTTg1), ns=0.0746 (HD-SaCas9 versus HD-SaCas9-HTTg1/gCas9), ns=0.9980 (HD-SaCas9-HTTg1 versus HD-SaCas9-HTTg1/gCas9). Two-way ANOVA with Tukey’s *post hoc* multiple comparisons test. (**E**) Improvement of coordination and balance ability in HD-SaCas9-HTTg1/gCas9 and HD-SaCas9-HTTg1 mice in rotarod test. N=15 mice per group; HD-SaCas9 versus HD-SaCas9-HTTg1/gCas9: \*\**p*=0.0065 (15 weeks), \*\*\**p*=0.0006 (16 weeks), \*\*\**p*<0.0001 (17-20 weeks). HD-SaCas9 versus HD-SaCas9-HTTg1/gCas9: ^###^*p*<0.0001 (15-20 weeks). Two-way ANOVA with Tukey’s *post hoc* multiple comparisons test. (**F**) Compared to HD-SaCas9, the survival time of HD-SaCas9-HTTg1/gCas9 mice was significantly extended and reached to a level consistent with that of HD-SaCas9-HTTg1 mice. *p*<0.0001 (HD-SaCas9 versus HD-SaCas9-HTTg1), *p*<0.0001 (HD-SaCas9 versus HD-SaCas9-HTTg1/gCas9), *p*=0.7174 (HD-SaCas9-HTTg1 versus HTTg1/gCas9). Non Tg-SaCas9 (N=8), HD-SaCas9 (N=10), HD-SaCas9-HTTg1 (N=10) and HD-SaCas9-HTTg1/gCas9 (N=9) mice. Log-rank (Mantel-Cox) test. Data were shown as mean ± SEM.

Transcriptomic dysregulation is a hallmark in HD patients and animal models (*52*). We investigated transcriptomes and found that compared to non-transgenic mice, BAC226Q had signature changes that were partially reversed by conventional and self-inactivating gene editing (**fig. S10A**). GO enrichment analysis was performed on the differentially expressed genes in BAC226Q-SaCas9 control, BAC226Q-SaCas9-HTTg1 and BAC226Q-SaCas9-HTTg1/gCas9 groups in comparison with Non tg-SaCas9 mice. Both the conventional CRISPR/Cas9 and the self-inactivating systems showed comparable efficacies in ameliorating transcriptomic-level dysregulation. The number and expression levels of genes involved in important pathways were significantly improved in gene-edited mice. Noticeably, most prominent changes were enriched in genes for postsynaptic structure and function, ion channel activity and transmembrane transporters (**fig. S10B**).

Therefore, we succeeded in developing a self-inactivating system that eliminated m*HTT* gene and protein, rescued deficits in motor functions and lifespan with the same high efficiency as the conventional CRISPR/Cas9, and that abolished Cas9 after a transient expression. This novel system could be highly beneficial for clinical applications.

### On and off-target effects of conventional and self-inactivating CRISPR/Cas9

BAC226Q contains human m*HTT* transgene and endogenous mouse *Htt* gene. To investigate the long-term cleavage effects on human m*HTT* gene, mouse brain genomic DNA was extracted at 11 months. Compared with control BAC226Q mice, the human m*HTT* gene sequence in BAC226Q-SaCas9-HTTg1 had been edited exactly from 3 bp upstream of PAM (**Fig. 7A**), consistent with previous findings (*53*). To compare the specificity of self-inactivating and conventional CRISPR systems, we applied primer-extension-mediated sequencing (PEM-seq), a method for detecting editing efficiency and off-target hotspots simultaneously because it can identify CRISPR/Cas9 editing events including indels, large deletions as well as genome-wide translocations with high sensitivity (*54*). In human *HTT* on-target site, the editing events were predominantly small indels, about 90% of which resulted in frameshift and premature stop codon (**Fig. 7, B and C**). In both strategies, no off-target events were found except for the predictable mouse *Htt* gene, which shares 20/21 sequence homology with human *HTT* gene target site (**Fig. 7D**). Even with 20/21 sequence homology, a 95% identical base pairs, HTTg1 only yielded off-target translocations with very low frequencies of 0.012% ± 0.001% in 2-month and 0.017% ± 0.004% in 12-month-old BAC226Q mouse brain striatum. HTTg1/gCas9 further significantly reduced HTTg1-induced off-target effects in 12-month-old mouse by 2.5-fold (**Fig. 7E**). For more clinical relevance, the off-target effect for HTTg1 was also investigated in human cell lines. In human iPSCs, neither HTTg1 nor HTTg1/gCas9 induced any off-target mismatches (**Fig. 7F**). In HEK293T cells, HTTg1 resulted only one off-target editing in chromosome 1 22,620,422-22,620,442 locus. Importantly, the self-inactivating HTTg1/gCas9 completely eliminated all off-target editing (**Fig. 7, G and H**). It is worth noting that gCas9 *per se* did not induce any off-target effect in mouse brain and human cell line. Taken together, these data confirmed the successful editing by SaCas9-HTTg1 and SaCas9-HTTg1/gCas9 on human *HTT* gene, and suggested that self-inactivating CRISPR/Cas9 may eliminate off-target effects and thus provide maximal safety for long-term therapeutic use.

**Fig. 7.**
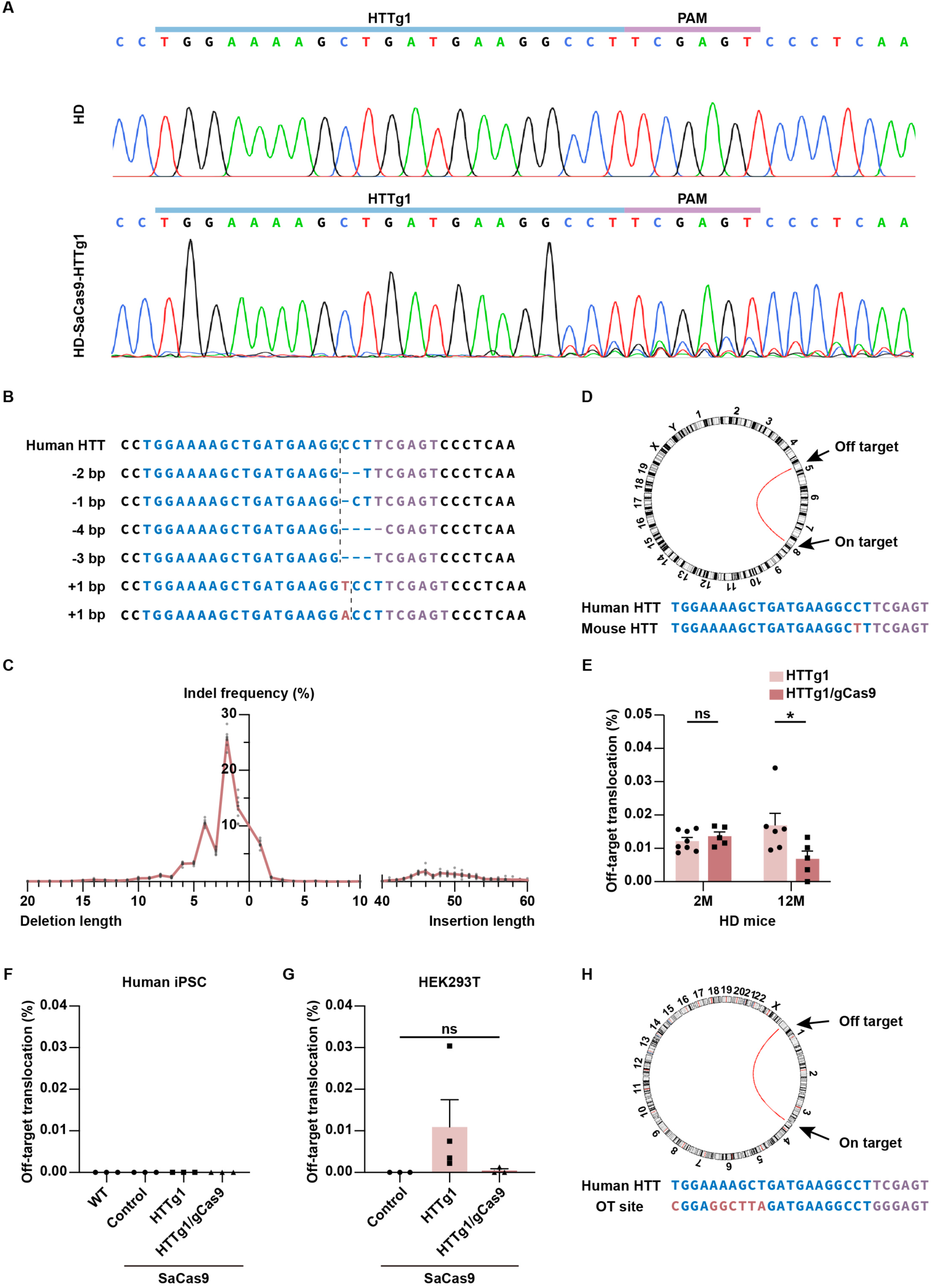
High gene editing specificity of HTTg1 and HTTg1/gCas9 in BAC226Q mouse model and human cell lines. (**A**) Sanger sequencing of human *HTT* exon 1 amplicons in control HD mouse (upper) and HD-SaCas9-HTTg1 mouse (lower). The genomic DNA editing by HTTg1 was demonstrated by the scrambled nucleotide signals in the lower panel. (**B** to **H**) Gene editing specificity detected by PEM-seq. (B and C) Indel pattens (B) and frequency (C) on human *HTT* in 12-month-old BAC226Q mice. Blue, guide RNA target site; Purple, PAM sequence; Red, insertions; Dash blue line, deletions. N=9 mice. (D) Circos plot for off-target site in BAC226Q mouse. Arrows indicate the on-target site of human *HTT* gene and the only off-target site of mouse *Htt* gene. The curved line connects the on-target and off-target site. Blue, guide RNA target site; Purple, PAM sequence; Red, mismatch. (E) Low frequencies calculated as off-target translocations over total editing events in the striatum of 2-month- and 12-month-old BAC226Q mice. N=8 for HD-SaCas9-HTTg1 (2 months), N=5 for HD-SaCas9-HTTg1/gCas9 (2 and 12 months) and N=6 for HD-SaCas9-HTTg1 (12 months); \**p*=0.0135; Two-way ANOVA with Bonferroni’s *post hoc* multiple comparisons test. (F) No off-target mismatch calculated as off-target translocations over total editing events in human iPSCs. N=3. (G and H) Off-target frequencies calculated as off-target translocations over total editing events (G) and circos plot (H) in HEK293T cells. Arrows indicate the on-target site of human *HTT* gene and off-target site on chromosome 1. The curved line connects the on-target and off-target site. Blue, guide RNA target site; Purple, PAM sequence; Red, mismatch. N=3 for SaCas9 and HTTg1/gCas9, N=4 for HTTg1; \*\*\**p*<0.0001; One-way ANOVA with Tukey’s *post hoc* multiple comparisons test. Data were shown as mean ± SEM.

## DISCUSSION

The revolutionary gene editing tools provide the possibility of root-cause cure for genetic disorders, especially monogenic and gain-of-toxic-function diseases, by eliminating or correcting mutated genes. To achieve clinical success, several aspects have to be satisfied, including high efficiency gene editing, effective delivery of gene editing tools to the neurons in pertinent brain structures, selection of intervention time points, elimination of off-target effects, etc. Our study strives to reach these goals with immediate clinical relevance.

Highly efficient gene editing is pivotal for disease rescue. In this study, we developed conventional and self-inactivating CRISPR gene editing systems, both of which successfully eliminated human m*HTT* gene, mHTT protein and aggregation in BAC226Q HD mouse brain. This was achieved by injection coverage of about 90% of striatal and 60% of M1 cortical area. In the injected areas, AAV9 transduced above 70% of neurons, among which 95% went through successful gene editing, leading to 60-90% mHTT protein and 90% of mHTT aggregation reduction. This mHTT suppression rescued deficient phenotypes in HD mouse model.

In HD patients and animal models, there is an age dependent and progressive mHTT accumulation of cytoplasmic aggregates and nuclear inclusions (*50*). This is well-recapitulated in BAC226Q mice. In our early injection experiment, cytoplasmic aggregates were greatly reduced and nuclear aggregates were almost eliminated. In later injection, the reduction of mHTT is still significant, although to a lesser degree. Importantly, injection at 4 months when BAC226Q start to show obvious cytoplasmic and nuclear mHTT aggregation, the deletion of m*HTT* gene not only prevented its expression, but also allowed neurons to gradually degrade the existing mHTT aggregates.

The primary neuropathology of HD occurs in the striatum and cerebral cortex (*55*), suggesting that we must take striatum, cortex and their interaction into account when we develop effective therapeutic strategy through directly targeting HTT (*56*). Our results illustrated that striatal and cortical disruption of human mutant HTT expression in BAC226Q mice by CRISPR/Cas9 was sufficient to slow the disease progression at multiple disease timepoints. This result is also the basis for choosing direct local injection into striatum and cortex as the method of delivery. The direct injection method circumvents issues such as considerable dilution of AAV in intravenous or intrathecal delivery (*57*), challenges in crossing the blood-brain barrier (*58*), transduction of brain areas irrelevant to HD pathogenesis, high AAV dosage induced toxicities (*59, 60*), immune responses (*61*) and off-target effects.

In neurodegenerative diseases including HD, glia-neuron interaction has been actively studied (*62–65*). In our study, AAV9 was stereotactically injected into mouse striatum and cortex, and transduced predominantly neurons, and much less glia. Interestingly, we found that CRISPR/Cas9-mediated *HTT* gene deletion in neuronal cells was adequate to rescue mutant phenotypes in BAC226Q mouse.

Since HD patients can be in presymptomatic, onset or symptomatic stages, the proper timing of gene editing intervention is important. Ideally, earlier m*HTT* gene disruption would yield better rescue benefits. However, how late can gene editing be performed and still be effective? Herein, we disrupted mHTT expression in BAC226Q mice at different time points: one month old BAC226Q mice before phenotypical onset; four months old BAC226Q mice of initial phenotypical onset of movement abnormalities; seven months old BAC226Q mice with severe mutant phenotypes. We found that as expected, earlier gene editing intervention has stronger therapeutic effects. However, it is exciting that gene editing at much later stages can still significantly extend their survival time. This systematic investigation on *HTT* gene disruption time points provides useful information on potential initiation time of treatments. Currently, HD clinical trials have included early symptomatic patients, and there is an ongoing discussion whether treatments can start at Stage 1 when there are biomarker changes but not overt motor symptoms (*66*).

In all clinical interventions including that for HD, a serious concern is that prolonged expression of SaCas9 may induce immune response and off-target effects (*67–71*). To address these concerns, we designed the self-inactivating CRISPR/Cas9 system where another sgRNA was introduced to delete SaCas9 in a short period of time. Comparable to the conventional CRISPR/Cas9, this self-inactivating system effectively deleted *HTT* gene, dramatically decreased mutant HTT aggregation and successfully ameliorated deficits in BAC226Q mice. Meanwhile, the self-inactivating system diminished the very low off-target frequency in conventional CRISPR. It is worth noting that the self-inactivating CRISPR/Cas9 could be a universal tool for maximal safety in clinical treatments.

In summary, our study demonstrated a clinically relevant gene editing strategy to efficiently ameliorate neuropathological phenotypes and behavioral deficits in a human genomic transgenic mouse model of HD. We illustrated that both conventional and self-inactivating CRISPR/Cas9 can efficiently and permanently delete mutant *HTT* gene *in vitro* and *in vivo*. This significantly reduced pathogenic protein aggregation, attenuated transcriptional dysregulation, improved motor performance, alleviated weight loss, and extended the lifespan of HD mice. It should be noted that these robust rescue effects were achieved by eliminating human m*HTT* with the present of wildtype mouse *htt*. Because of this and that mouse models may differ from higher animals, for clinical trial readiness, we are testing our system in cynomolgus macaque for safety, gene editing efficiency and AAV dosage. We hope that our CRISPR/Cas9-mediated gene editing via AAV delivery is feasible and safe for HD and a large spectrum of hereditary neurological disorders.

## MATERIALS AND METHODS

### Study design

The aim of this study was to develop a CRISPR-based gene editing as a therapy for HD with maximal safety. This study investigated the long-term rescue and off-target effects in a robust HD mouse model BAC226Q at different time points of intervention. Given the high HTT editing efficiency of CRISPR in HEK293T cell line, we therefore packaged our editing system in AAV9 and delivered it to the striatum and cortex of BAC226Q HD mice, evaluating the *in vivo* editing effects at different post-injection time points by sequencing, Western-blot, immunostaining and behavioral tests. The dosage of AAV vectors was determined based on published papers and our pilot studies. The cell line used in this study was indicated in the place of related results shown in the text, figure legends, and Materials and Methods. Animal size was determined on the basis of the previous studies in BAC226Q mouse. Experimental endpoints were determined by cardinal parameters of age-dependent pathogenesis in BAC226Q. Both sexes were used in Western-blot and staining studies and only male mice were used in behavioral and sequencing studies. Animals were randomly assigned to the control and experimental groups, and researchers were blinded to genotype and treatment information during the experiment but not during analysis. Details on the use of cell line and animal in different experiments could be found in the corresponding sections of Materials and Methods. Biological replicates were indicated by N values in the figure legends. No data were excluded.

### Statistical analysis

Unless otherwise mentioned, one-way analysis of variance (ANOVA) with Tukey’s *post hoc* multiple comparisons test or two-way ANOVA with Tukey’s *post hoc* multiple comparisons test were used for statistical comparison. Survival data were analyzed by Log-rank (Mantel Cox) testing. Results were expressed as a mean ± standard error of the mean (SEM). Statistical tests were performed and graphs were generated using GraphPad Prism (GraphPad Software, Inc.). Statistical significance was defined as *p*<0.05. Statistical significance level was set as follows: \**p*<0.05, \*\**p*< 0.01, \*\*\**p*< 0.001.

## Acknowledgements

We thank Prof. Hongkui Deng at Peking University for the supply of hCiPSC; Prof. Xiaowei Chen and Dr. Xiao Wang at Peking University for technical support in AAV virus production and purification; Prof. Gillian Bates at UCL Queen Square Institute of Neurology for the supply of the S830 antibody; Prof. Xiaojiang Li at Jinan University for the supply of the EGFP-hHTT-20Q/120Q constructs. The authors thank the Laboratory Animal Center and staff members for their assistance in animal care and maintenance. The authors thank National Center for Protein Sciences at Peking University in Beijing, China, for assistance with the behavioral experiments and Dr. Yonglu Tian for technical help.

## Author Contributions

Conceptualization: S.Z., C.L.

Methodology: Y.Dai, Z.A., Y.Ding, W.H., J.Y., L.O., J.H.

Investigation: Y.Dai, Z.A., Y.Ding, W.H. J.Y., L.O.

Funding acquisition: C.L.

Project administration: S.Z.

Supervision: C.L., S.Z.

Writing – original draft: Y.Dai, Z.A., S.Z., C.L.

Writing – review & editing: Y.Dai, Z.A., S.Z., C.L.

## Data and materials availability

All data are available in the main text or the supplementary materials.

## Supplementary Materials

### Materials and Methods

#### Molecular cloning

A single vector AAV-SaCas9 system containing Cas9 from *Staphylococcus aureus* (SaCas9) and its sgRNA scaffold was obtained from Addgene (plasmid #61591). sgRNAs targeting human *HTT* and SaCas9 were designed based on PAM sequence (5’-NNGRRT-3’) and by CRISPR RGEN Tools (http://www.rgenome.net/). sgRNA sequences targeting human *HTT* are as follows: HTTg1: 5’-TGG AAA AGC TGA TGA AGG CCT-3’; HTTg2: 5’-GAA AAG CTG ATG AAG GCC TTC-3’. AAV-EGFP-gCas9 or AAV-mCherry-gCas9 which contain EGFP or mCherry were constructed by replacing SaCas9 on vector AAV-SaCas9. sgRNAs targeting SaCas9 are listed in table S1.

Cas9 was replaced by EGFP or mCherry in the AAV vector backbone via *Age*I and *Eco*RI enzyme digestion to generate AAV-EGFP and AAV-mCherry vectors, respectively. Top and bottom strands of oligos for each sgRNA were phosphorylated and annealed into duplex. HTTg1 and HTTg2 were inserted into AAV-SaCas9 vector via *Bsa*I enzyme digestion. Similarly, gCas9 was cloned into the AAV-EGFP and AAV-mCherry vector through *Bsa*I digestion. The ligation products were transformed into Stbl3 chemically competent cells (TransGen, CD521) and plasmid DNA was extracted from cultures using the QIAprep spin miniprep kit (QIAGEN, 27104) according to the manufacturer’s instructions. sgRNA sequences were verified by Sanger sequencing using the LKO.1 5’ primer: 5’-GAC TAT CAT ATG CTT ACC GT-3’. EGFP and mCherry sequences were verified using the EGFP-forward primer: 5’-CGA AGG CTA CGT CCA GGA GC-3’ and the mCherry-forward primer: 5’-ACA ACC GGT ATG GTG AGC AAG-3’, respectively. AAV plasmids were then extracted by EndoFree plasmid maxi kit (QIAGEN, 12362) for subsequent virus production and purification.

#### Culture and transfection of HEK293T cell

Human embryonic kidney (HEK) 293T cells (ATCC, CRL-1573) were cultured in Dulbecco’s modified Eagle’s medium (DMEM, Cytiva, SH3024301) supplemented with 10% (v/v) fetal bovine serum (FBS, Thermo Fisher), 1% (v/v) penicillin-streptomycin (Gibco, 21103049) and 1% (v/v) L-glutamine (Gibco, 25030081) in a humidified 5% CO_2_ atmosphere at 37°C. Cells were transfected with 2.5 μg AAV-SaCas9-sgRNA vectors and 2.5 μg pEGFPc3-human HTT exon 1-20 CAG/pEGFPc3-human HTT exon 1-120 CAG (EGFP-hHTT-20Q/120Q) at 70% confluency with lipofectamine 2000 (Invitrogen, 11668030); In self-inactivating system, cells were co-transfected with 0.8 μg AAV-SaCas9-HTTg1/AAV-SaCas9, 0.8 μg AAV-mCherry-gCas9/AAV-mCherry and 0.8 μg EGFP-hHTT-20Q/120Q. For *in vitro* validation of cleavage effects, EGFP fluorescence was imaged under microscope (Carl Zeiss, Vert.A1) 24 or 48 h after transfection, and then protein was harvested for Western-blot. For PEM-seq assay, cells were transfected with 5 μg AAV-SaCas9-HTTg1/AAV-SaCas9 and 5 μg AAV-mCherry-gCas9/AAV-mCherry, then mCherry-positive cells were sorted by flow cytometry (BeckMan Coulter, Astrios EQ) 24 h after transfection, followed by genomic DNA extraction.

#### Culture and nucleofection of human iPSC

Human chemical induced pluripotent stem (hCiPSC) cell line (#1217-2) was obtained from Prof. Hongkui Deng at Peking University. hCiPS cells were cultured on Matrigel (Corning, 354230) coated plates in mTeSR^TM^ Plus Medium (STEMCELL, 100-0276) in a humidified 5% CO_2_ atmosphere at 37°C. When reached to 80% confluence, cells were dissociated with ReLeSR^TM^ (STEMCELL, 05872) and passaged with a 1:10 split ratio. Detached cell aggregates were first cultured in mTeSR^TM^ Plus Medium supplemented with 10 μM Y-27632, and replaced by fresh mTeSR^TM^ Plus Medium the next day. Culture medium were changed every one or two days. hCiPS cells were transfected by nucleofection with Lonza P3 primary cell kit under CM115 program (Lonza). For each 6-well, 2.5 μg of AAV-SaCas9-HTTg1/AAV-SaCas9 vectors and 2.5 μg of AAV-mCherry-gCas9/ AAV-mCherry were transfected. For PEM-seq assay, mCherry-positive hCiPS cells were sorted by flow cytometry 48 h after nucleofection. Cells were cultured in mTeSR^TM^ Plus Medium supplemented with 10 μM Y-27632 for 24 h after nucleofection and sorting, and replaced by fresh mTeSR^TM^ Plus Medium for further culturing.

#### Mice

Mice were housed in the specific pathogen-free (SFP) facility at Peking University in accordance with guidelines. The room was controlled with 12 h light/dark cycle, 40-60% humidity and 22 ± 1°C temperature. Food and water were always accessible. All mice used in this research were FVB/N (Taconic) background, as BAC226Q mice in this background showed the most robust HD-like phenotypes. BAC226Q mice were bred with wild-type non-transgenic mice and confirmed by genotyping which produced a 403 bp PCR fragment (Forward primer: 5’-GTA TAT GCT GCT GCC TGC AA-3’; reverse primer: 5’-AGG GGA CAG TGT TGG TCA AG-3’). All mice experimental procedures were approved by the IACUC of Peking University.

#### Western-blot

Mouse brains were dissected in ice-cold PBS. Lysates were prepared by ultrasonication in RIPA buffer (150 mM NaCl, 0.1% SDS, 1% Sodium deoxycholate, 50 mM Triethanolamine, 1% NP-40, pH 7.4, supplemented with Caspase inhibitor Boc-D-FMK (Abcam, ab142036) and Protease inhibitor cocktail (Sigma, 111M4009)) using a sonic dismembrator (Fisherbrand, FB120), pulsed on for 1 sec and pulsed off for 2 sec, repeated for 20-25 times at 20% amplitude on ice. The lysates were incubated at 4°C for 60 min, followed by centrifugation at 4°C for 20 min at 16,100 *g* to remove insoluble components. Protein concentration was measured by BCA protein assay using Pierce BCA Protein Assay Kit (Thermo Scientific, 23225) and the absorbance was detected at 562 nm. Protein samples were prepared for loading by heating in NuPAGE 4×LDS sample buffer plus 10×sample reducing agent (Invitrogen) according to the manufacturer’s protocol and heated for 10 min at 70°C. A total amount of 90-120 μg of protein was loaded onto 4%-12% NuPAGE Bis-Tris gels (Invitrogen, NP0321/NP0336) using MOPS running buffer, and wet-transferred to an Immobilon-FL PVDF membrane (Millipore, IPFL00010). Lysates from HEK293T cells were prepared in Triton X-100 buffer (150 mM NaCl, 50 mM Tris-HCl, pH 7.4, 1 mM EDTA, 1% Triton X-100). The lysates were incubated at 4°C for 30 min, followed by centrifugation at 4°C for 20 min at 16,100 *g* to remove insoluble components. Protein concentration was measured by BCA protein assay. Protein samples were prepared for loading by heating in 4×SDS sample buffer and heated for 10 min at 95°C. A total amount of 30 μg of protein was loaded onto SDS-PAGE gels using running buffer, and wet-transferred to an Immobilon-FL PVDF membrane. Blots were incubated with Odyssey blocking buffer (LI-COR, 927-40000) for 60 min. After washing with TBST, blots were incubated with primary antibodies diluted in blocking buffer at 4°C overnight. Blots were washed 3 times (10 min each) and incubated with fluorescently labeled IRDye 680RD goat anti-mouse (1:10,000, LI-COR, 926-68070) or goat anti-rabbit (1:10,000, LI-COR, 926-68071) secondary antibodies in TBST for 60 min at room temperature. Protein was detected in the 700 nm channel using Odyssey CLx imager (LI-COR). Following primary antibodies were used: 1C2 (1:5,000, mouse, Millipore, MAB1574), GFP (1:8,000, rabbit, Abcam, ab183734), HA (1:2,000, mouse, Abcam, ab18181) and β-actin (1:2,000, rabbit, Cell Signaling, 4970).

#### Immunofluorescent staining

Mice were anesthetized, perfused with fresh 4% paraformaldehyde in PBS, and post-fixed overnight in the same fixative solution. Fixed brains were sliced at 40 μm thickness with a vibrating blade microtome (Leica, VT1200S). The brain slices were incubated in PBS supplemented with 0.3% (v/v) Triton X-100 for 30 min in room temperature and then blocked with 10% (v/v) goat serum (ZSGB-BIO, ZLI-9021) in PBS supplemented with 0.1% or 0.3% Triton X-100 for 60 min at room temperature. Following incubation of brain slices with primary antibodies at 4°C overnight and washes with PBS supplemented with 0.1% Triton X-100, fluor-conjugated secondary antibodies (anti-mouse, anti-rabbit or anti-sheep Alexa Fluor 488, 555, 594 or 647, 1:1,000, Thermo Fisher Scientific) were added to the samples for 60 min at room temperature. Sections were cover slipped by antifade mountant with Hoechst 33342 nuclear stain (Thermo Fisher Scientific, P36985). Images were taken by Zeiss LSM 780 or LSM 880 confocal microscope (Carl Zeiss), whole brain section images were taken by Axio Scan Z1 (Carl Zeiss). Following primary antibodies were used: mEM48 (1:1,000, mouse, Millipore, MAB5374), S830 (1:5,000, sheep, kindly provided by Professor Gillian Bates from University College London), HA (1:1,000, rabbit, Cell Signaling Technology, 3724), HA (1:1,000, mouse, Abcam, ab18181), GFAP (1:1,000, rabbit, Abcam, ab7260), NeuN (1:1,000, mouse, Abcam, ab104224) and DARPP-32 (1:500, rabbit, Abcam, ab40801).

#### Immunohistochemistry

Mouse brain sections were prepared by the same procedure as the immunofluorescent staining samples. Sections were stained according to instructions provided with the ABC and DAB substrate kits (Vector Labs). Briefly, Sections were first processed free-floating with blocking serum and primary antibodies. After treatment with biotinylated secondary antibody, followed by avidin-biotinylated horseradish peroxidase complexes and incubation with diaminobenzidine, sections were mounted onto slides. And brain sections were dehydrated using the set up as 30% alcohol, 50% alcohol, 70% alcohol, 95% alcohol, 100% alcohol-I, 100% alcohol-II (2 min each); xylene-I and xylene-II (10 min each). Then sections were cover slipped by DPX mounting medium (Sigma, 06522) and imaged under Axio Scan Z1 microscope. Following primary antibodies were used: S830 (1:25,000, sheep, kindly provided by Professor Gillian Bates from University College London), HA (1:1,000, rabbit, Cell Signaling Technology, 3724) and GFAP (1:2,000, rabbit, Abcam, ab7260).

#### AAV vector production

HEK293T cells were plated on ten 15-cm dishes and transfected at 90% confluence by adding 5 mL of the plasmid and linear polyethylenimine (PEI, Sigma-Aldrich, 765090) mixture to each plate. The mixture was prepared with 70 μg AAV9 vector (AAV-SaCas9-HTTg1, AAV-SaCas9, AAV-EGFP-gCas9 or AAV-EGFP), 200 μg Ad-Helper plasmid, 70 μg AAV-Rep/Cap plasmid and PEI (1 μg/μL), the PEI to DNA ratio was 5:1 (v/g) in our experiment. The mixture was filled to 50 mL with DMEM and mixed well. Cells were collected with culture medium 60 h after transfection and centrifuged at 1,000 rpm for 10 min. Cell pellet was re-suspended in 5 mL cell lysis buffer (150 mM NaCl, 20 mM Tris, pH 8.0).

Freeze-thaw the cell lysate for 3 times by changing between lipid nitrogen and 37°C water bath to release virus from the destroyed cell membrane. Cell lysate was added with MgCl_2_ and Benzonase (Sigma, E8263-25k) to a final concentration of 1 mM and 250 U/mL respectively, followed by incubation at 37°C for 15 min to dissolve the DNA/protein aggregation. The supernatant was collected after centrifugation of the cell lysate at 4,000 rpm for 30 min at 4°C.

The viral solution was loaded on the iodixanol gradient solution for purification, the gradient was prepared in the following order (from top to bottom): 6 mL of 17% iodixanol (5 mL 10×PBS, 0.05 mL 1 M MgCl_2_, 0.125 mL 1 M KCl, 10 mL 5 M NaCl, 12.5 mL Optiprep (Sigma, D1556) and H_2_O up to 50 mL), 6 mL of 25% iodixanol (5 mL 10×PBS, 0.05 mL 1 M MgCl_2_, 0.125 mL 1 M KCl, 20 mL Optiprep and H_2_O up to 50 mL), 5 mL of 40% iodixanol (5 mL 10×PBS, 0.05 mL 1 M MgCl_2_, 0.125 mL 1 M KCl, 33.3 mL Optiprep and H_2_O up to 50 mL) and 4 mL of 60% iodixanol (0.05 mL 1 M MgCl_2_, 0.125 mL 1 M KCl, 50 mL Optiprep). The sample was centrifuged at 53,000 rpm for 160 min at 14°C and the viral fraction was harvested in the 40% layer with a syringe.

The viral fraction was mixed with PBS solution and 10% Poloxamer 188 solution (F188) (1:10,000, Sigma, P5556) and then it run through the Amacon 100K filter (Millipore Sigma, UFC910008) at 3,500 rpm for 20 min at 4°C to remove the iodixanol and concentrate the virus. The filtrate was removed, and the viral fraction was again filled with F188-containing PBS solution. The filter was centrifuged at 3,500 rpm for 20 min at 4°C. The filtrate was discarded again, and the viral fraction was mixed with PBS. The virus was concentrated by centrifugation at 3,500 rmp for 20 min at 4°C until the viral volume was about 500 μL.

Virus titer was measured by quantitative PCR. For AAV-SaCas9-HTTg1 and AAV-SaCas9, common U6-forward primer (5’-CGA TAC AAG GCT GTT AGA GAG-3’) was used, with reverse primers were AAV-HTTg1-Reverse (5’-AGG CCT TCA TCA GCT TTT CCA-3’) and AAV-SaCas9-Reverse (5’-GTG GTC TCC GGT GTT TCG TCC-3’) respectively. For AAV-EGFP-gCas9 and AAV-EGFP, the primers used are AAV-EGFP-Forward (5’-CGA AGG CTA CGT CCA GGA GC-3’) and AAV-EGFP-Reverse (5’-CGA TGT TGT GGC GGA TCT TG-3’). SDS-PAGE followed by Coomassie blue staining was performed to check the purity of the virus vectors, in which the only proteins visible in the gel were the VP1, VP2 and VP3 proteins that make up the capsid particle, migrating at approximately 87 kDa, 73 kDa and 62 kDa, respectively.

#### Stereotaxic injection procedure

Stereotaxic administration of AAV9 were performed on 1, 2, 4 or 7-month-old mice. Mice were anesthetized with 1.5% isoflurane inhalation and stabilized in a stereotaxic instrument (RWD, 68019). Anterior-posterior (AP), medial-lateral (ML) and dorsal-ventral (DV) injection coordinates were calculated relative to the bregma. The injection coordinates for striatum were as follows: AP = 0.8 mm, ML = ±2.1 mm, and DV = −3.1 mm (1 and 2-month-old mice); AP = 0.7 mm, ML = ±2.0 mm, and DV = −3.1 mm (4-month-old mice); AP = 0.6 mm, ML = ±2.1 mm, and DV = −3.1 mm (7-month-old mice). The injection coordinates for cortex were AP = 1.5 mm, ML = ±1.5 mm, and DV = −1.0 mm (mice of all ages). AAV9 was bilaterally injected into the striatum (2.5 μL) and primary motor cortex (0.5 μL) of mice. In conventional CRISPR/Cas9 system, a total dose of 1.2×10^11^ viral genomes (vg) of AAV-SaCas9-HTTg1 or AAV-SaCas9 was injected into each mouse brain (virus titer: 2×10^13^ vg/mL). In self-inactivating system, AAV-EGFP-gCas9 or AAV-EGFP were co-injected with AAV-SaCas9-HTTg1 or AAV-SaCas9 at 1:1 molar ratio, and a total dose of 1.2×10^10^ vg of virus was administrated for each mouse (virus titer: 2×10^12^ vg/mL). Injection was performed through a microinjection pump (RWD, 788130)-connected Hamilton syringe at an injection rate of 0.3 μL/min in striaum and 0.1 μL/min in cortex. A 1701 Hamilton microsyringe (Hamilton, 7853-01) with 33-gauge needle (Hamilton, 7803-05) was used to deliver the virus. The needle was left in place for 15 min after each injection to minimize upward flow of viral solution after raising the needle. After surgery the mice were placed on a heated blanket to recover from the anesthetic.

#### Mouse behavioral analysis

All behavioral tests were performed during the dark phase of the light cycle and at the same time of day. Mice were handled under the same conditions by one investigator who was blind to the genotype and treatment of the mice.

##### Rotarod test

Rota-rod tests were operated on a rotarod treadmill (Med Associates, Inc., ENV-574M). Mice were trained at a constant speed of 10 rpm/min for 1 min per trial for 3 trials per day for 3 consecutive days. During the training trials, mice that fell were gently returned to the rotarod treadmill. Animals were tested for 3 trials per day for 3 consecutive days in every week, allowing at least 15 min of rest between each trial. Mice were tested with the rotarod accelerating from 5 to 40 rpm with a maximum period of 300 sec. The latency to fall was recorded and the mean of all trials was used for statistical analysis. The latency to fall was defined by either the animal falling from the rotarod or when the animal holds the rod for more than 3 cycles.

##### Open-field

The open field consisted of a clear glass box with evenly distributed photobeams (27.3×27.3×20.3 cm^3^, Med Associates, Inc., ENV-510). Mice were placed in the open field over a 10 min period and kept track of locomotion. For each time points, mice were tested for 3 trials per day for 3 consecutive days. Average ambulatory distance or total travel distance were recorded, and the mean of all trials was used for statistical analysis. Representative figures of heatmaps showing the time and location to stay in the open field box were drawn by R (https://www.r-project.org/), referring to the average of 9 sessions for one mouse.

##### Catwalk

Gait of mice was analyzed with CatWalk XT (Noldus Information Technology, Wageningen, Netherlands). The CatWalk system consists of an enclosed walkway on a glass plate that is traversed by a mouse from one side of the walkway to the other. A completely internally reflected green light can emit only at areas where the paws make contact with the glass plate. It scatters and illuminates the paw prints. Paw prints are captured by a highspeed video camera that is positioned underneath the walkway. The video images with paw prints are used in the footprint classification and further analysis. Mice were trained and adapted to walk across the 70 cm glass walkway in an unforced manner without any interruption or hesitation at least 4 times a day for at least 4 days prior to measurement. After training, mice footprints were measured 3 times per day on 3 consecutive days at each time point. Runs with any wall climbing, grooming, and staying on the walkway were not analyzed. Mice that failed the catwalk training were excluded from the study. An average number of 3-6 compliant trials made by each mouse were used for analysis. Analysis was performed using the CatWalk XT 10.6 Software. In catwalk performance, there are six step sequence patterns of walking for mice: AA (RF-RH-LF-LH), AB (LF-RH-RF-LH), CA (RF-LF-RH-LH), CB (LF-RF-LH-RH), RA (RF-LF-LH-RH) and RB (LF-RF-RH-LH). Step sequence regularity index is a measurement of walking coordination and regular step patterns (*72*).

##### Body weight

The body weight of mice was measured in grams every week after surgery.

##### Survival

The mice were kept in mix genotype with 3 to 5 mice per cage. After surgery, mice were monitored weekly for survival for at least 13 months.

#### PEM-seq

Genomic DNA of mouse brain and cell line samples were extracted by FastPure DNA isolation kit (Vazyme, DC102-01) following manufacturer’s instructions and were sonicated to 300-500 bp. And biotinylated 5’-CTC AGG TTC TGC TTT TAC CTG CG-3’ was used for primer extension followed by bridge adapter ligation. 5’-CCG AGG CCT CCG GGG ACT GC-3’ was used for nested PCR followed by tagged PCR with primers compatible for Illumina Hiseq sequencing. The raw data processing was described before (*54*). Editing efficiency was the percentage of indels and translocations to total events; and the off-target translocation was the percentage of junctions at off-target-site to the total editing events.

#### Sanger sequencing

The sequence of disrupted human HTT gene in BAC226Q mouse was characterized by Sanger sequencing. Mouse brain genomic DNA was extracted as previously described. 100 ng of genomic DNA was amplified with human *HTT*-specific primer (hsHTT forward: 5’-CTG CCG GGA CGG GTC CAA GAT-3’; hsHTT reverse: 5’-TGC AGC GGC TCC TCA GCC AC-3’) with the KAPA 2G Robust HotStart PCR kit (Roche, KE5507). The resulting PCR products of human *HTT* exon 1 fragments were purified with the QIAquick PCR purification kit (Qiagen, 28104) according to the manufacturer’s instructions and sequenced by Sanger sequencing.

#### RNA-seq and data analysis

The total stratal mRNA from 4-month-old Non Tg-SaCas9, BAC226Q-SaCas9, BAC226Q-SaCas9-HTTg1 and BAC226Q-SaCas9-HTTg1/gCas9 mice were isolated by FreeZol Reagent (Vazyme, R711-01). The RNAs were sent to Beijing Novogene Technology for transcriptome sequencing and analysis. 10 μg of total RNA for each qualified sample was used in the NEBNext Ultra RNA library prep kit (E7530L; NEB) to build the library. After cluster generation, the library preparations were sequenced on an Illumina Novaseq platform and 150 bp paired-end reads were generated. Index of the reference genome was built using Hisat2 v2.0.5 and paired-end clean reads were aligned to the reference genome using Hisat2 v2.0.5. Feature Counts v1.5.0-p3 was used to count the number of reads for each gene, and then the FPKM of each gene was calculated according to the length of the gene and reads count mapped to this gene. For differential gene analysis, the number of reads was normalized using DESeq software. The Benjamini and Hochberg’s method was used for adjusting the resulting *p*-values, and genes with absolute fold change>1 & padj <0.05 were considered differentially expressed genes.

### Supplementary figures

**Fig. S1.**
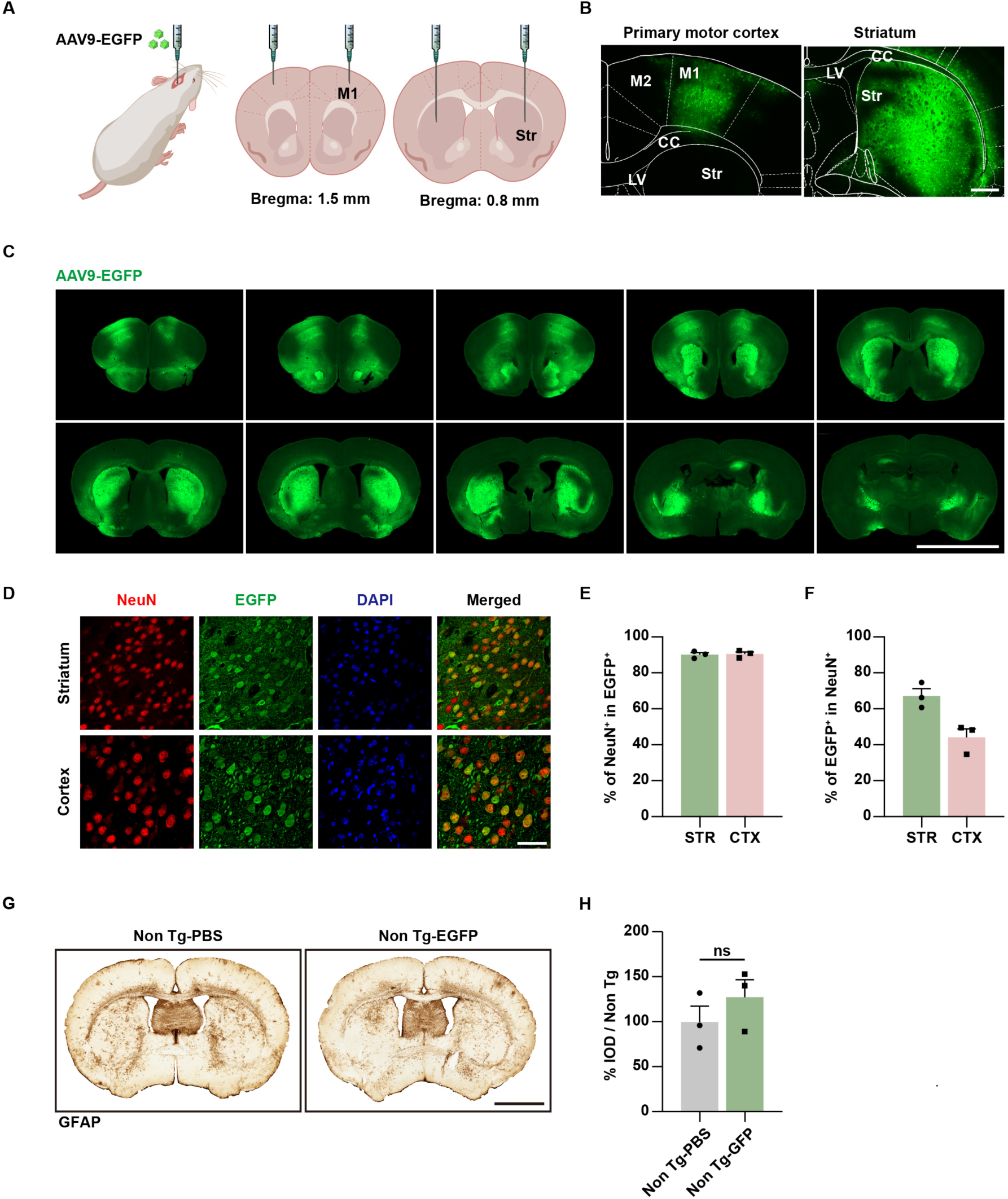
Highly efficient transduction of AAV9 in mouse brain without inducing immunoreactivity. (**A**) Schematic diagram of AAV vectors delivered by intraparenchymal injection. AAV9-EGFP was stereotactically injected into the striatum and primary motor cortex of 1-month-old mice. M1, primary motor cortex; Str, striatum. b,c, Efficient brain transduction of AAV9. (**B**) Transduction pattern of AAV9-EGFP in mouse primary motor cortex (left) and striatum (right). Images were taken one month after AAV9 injection. Scale bar: 500 μm. M2, secondary motor cortex; CC, corpus callosum; LV, lateral ventricle. (**C**) AAV9-EGFP indicated virus was transduced well throughout striatum and cortex. Sections were selected to imaging every 400 μm and images were taken ten months after AAV injection. Scale bar: 5000 μm. (**D** to **F**) High transduction efficiency of AAV9 in neurons 3 months post injection. (D) Immunofluorescent staining of mouse striatum (upper) and cortex (lower) for EGFP (green), NeuN (red) and DAPI (blue). Scale bar: 50 μm. Of all AAV9 transduced cells (EGFP^+^), neurons (NeuN^+^) were 90.14% ± 0.15% in striatum and 90.48% ± 1.19% in cortex (E). Of all neurons (NeuN^+^), 67.17% ± 4.04% in striatum and 44.10% ± 4.75% in cortex were transduced by AAV9 (EGFP^+^) (F). Note: the injection dosage was three times lower in cortex than in striatum. N=3 pairs of mice. (**G** and **H**) Unchanged reactive astrogliosis by AAV-EGFP. (G) GFAP signal was detected in 11-month-old mice injected with AAV-EGFP or PBS at 1 month. Scale bar: 2000 μm. (H) Quantification of GFAP integrated option density (IOD). *p*=0.7514. N=3 mice per group. Data were shown as mean ± SEM. One-way ANOVA with Tukey’s *post hoc* multiple comparisons test.

**Fig. S2.**
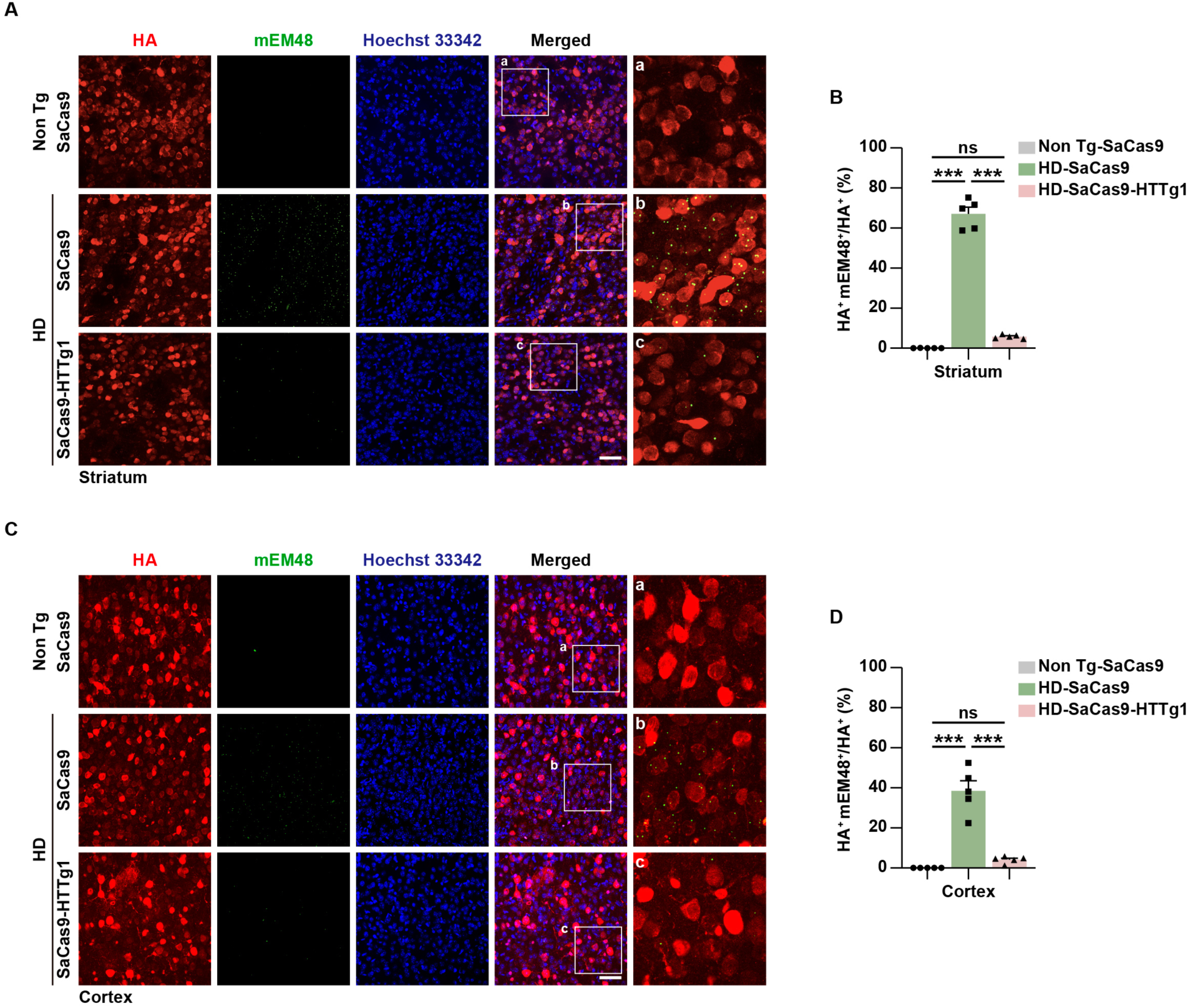
Decreased human mHTT by AAV-SaCas9-HTTg1 in BAC226Q mice. (**A** to **D**) Dramatic reduction of mEM48-positive mHTT aggregates (green) in HA-labeled SaCas9-positive cells (red) in the striatum (A and B) and primary motor cortex (C and D) of 7-month-old HD-SaCas9-HTTg1 mice. Enlarged figures were shown on the right. Scale bar: 50 μm. The percentage of mEM48-positive cells among SaCas9-HA expressing cells in striatum (B) and primary motor cortex (D). N=5 mice per group; \*\*\**p*<0.0001. Data were shown as mean ± SEM. One-way ANOVA with Tukey’s *post hoc* multiple comparisons test.

**Fig. S3.**
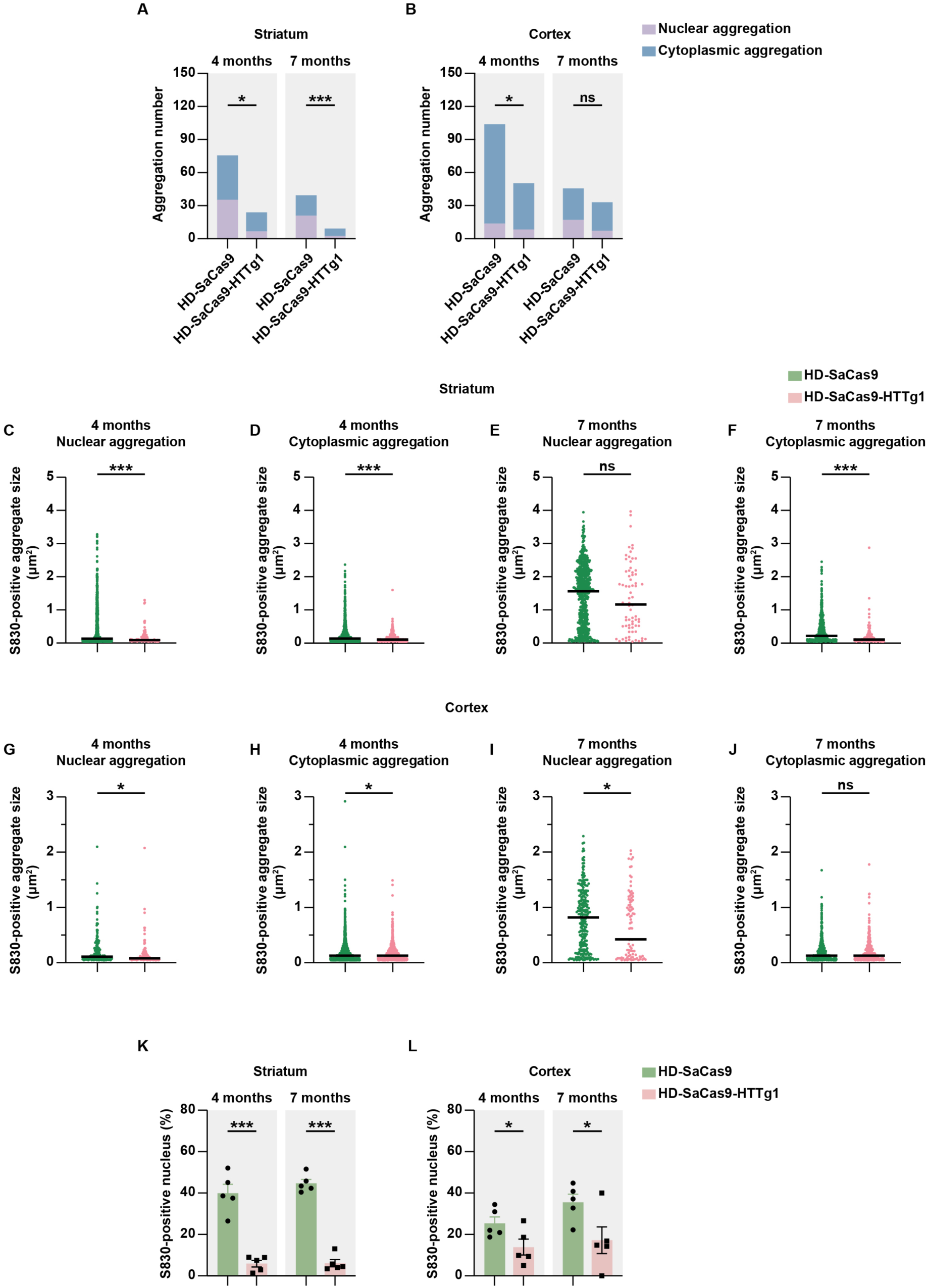
Reduction of nuclear and cytoplasmic mHTT aggregation by AAV-SaCas9-HTTg1 in BAC226Q mice. (**A** and **B**) Quantification of nuclear and cytoplasmic S830-positive mHTT aggregate number in HTT-g1 mouse striatum (A) and primary motor cortex (B). Striatum: \**p*=0.0176, \*\*\**p*<0.0001; Cortex: \**p*=0.0414, ns=0.1793. (**C** to **J**) Quantification of nuclear and cytoplasmic S830-positive mHTT aggregate size in HD-SaCas9-HTTg1 mouse striatum (C to F) and cortex (G to J). \*\*\**p*<0.0001 (C, D and F), ns=0.2817 (E), \**p*=0.0479 (G), \**p*=0.0485 (H), \**p*=0.0330 (I), ns=0.6053 (J). (**K** and **L**) Percentage of S830-positive nucleus in HD-SaCas9-HTTg1 mouse striatum (K) and cortex (L). \*\*\**p*<0.0001, \**p*=0.0472 (4 months), \**p*=0.0408 (7 months). N=5 mice per group. Data were shown as mean ± SEM. Unpaired two-tailed *t*-test.

**Fig. S4.**
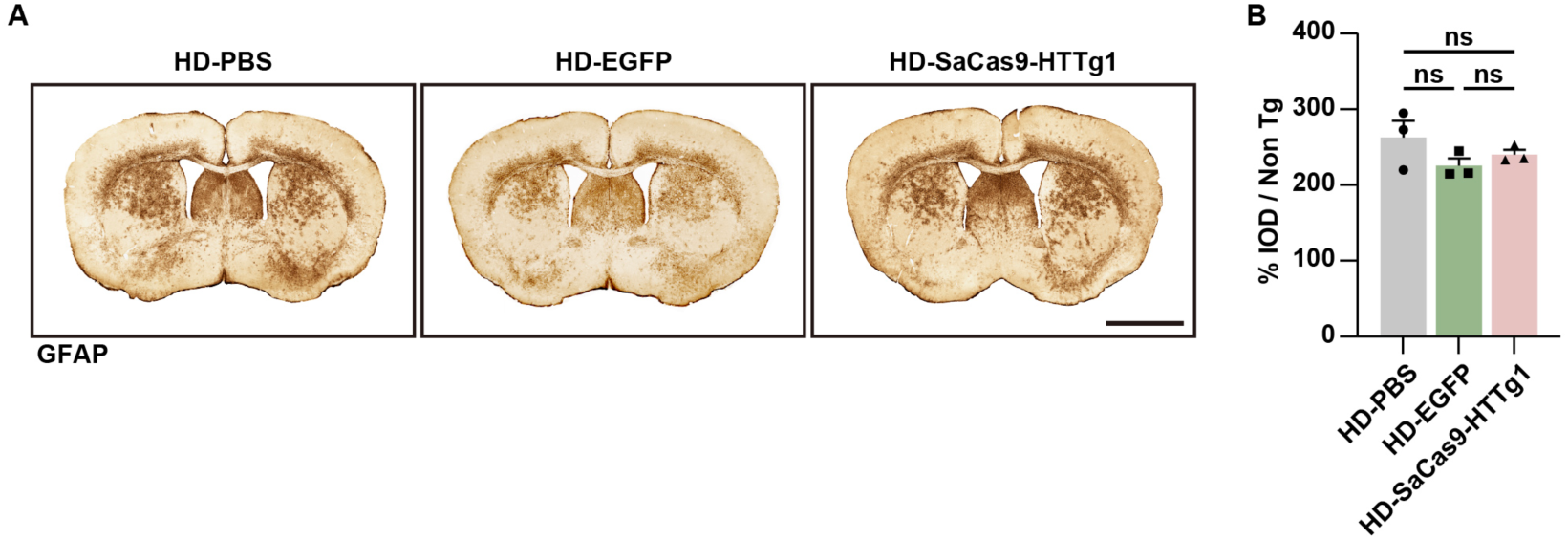
Unchanged reactive astrogliosis by AAV-SaCas9-HTTg1 in BAC226Q mice. (**A**) Unchanged GFAP signal in 11-month-old BAC226Q mice injected with AAV-SaCas9-HTTg1, AAV-EGFP or PBS. Scale bar: 2000 μm. (**B**) Quantification of GFAP integrated option density (IOD) in (A). No AAV9-EGFP treatment induced GFAP increase (HD-EGFP versus HD-PBS, *p*=0.5179), and AAV9-SaCas9-HTTg1 treatment did not change GFAP signal (HD-SaCas9-HTTg1 versus HD-EGFP, *p*=0.9626). N=3 mice per group. Data were shown as mean ± SEM. One-way ANOVA with Tukey’s *post hoc* multiple comparisons test.

**Fig. S5.**
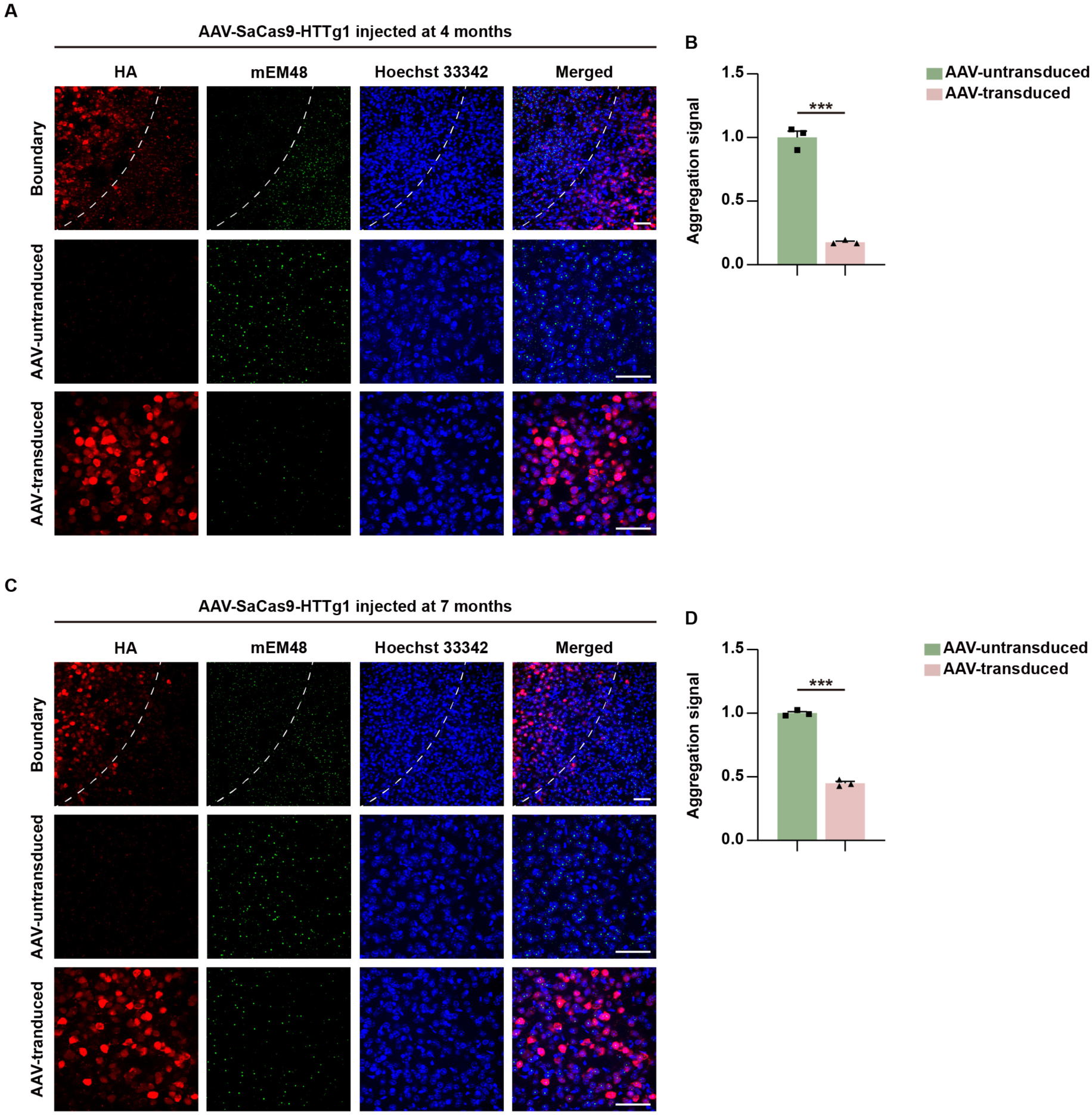
Significant human mHTT reduction by gene editing at 4 and 7 months with different efficiency. (**A** and **B**) Significant reduction of mEM48-positive mHTT aggregates (green) in the striatum at the age of 11 months from 4-month-injected HD mice. (A) The boundary (top panels) and representative figures of AAV9-SaCas9-HTTg1-untransduced (middle panels) and transduced area (bottom panels). (B) Quantification of mHTT aggregation signal in (A). (**C** and **D**) Significant reduction of mEM48-positive mHTT aggregates (green) in the striatum at the age of 11 months from 7-month-injected HD mice. (C) The boundary (top panels) and representative figures of AAV9-SaCas9-HTTg1-untransduced (middle panels) and transduced area (bottom panels). (D) Quantification of mHTT aggregation signal in (C). mHTT (green) and SaCas9 (red) were detected by mEM48 and anti-HA antibody, respectively. Scale bar: 50 μm. N=3 mice per group; \*\*\**p*<0.0001. Data were shown as mean ± SEM. Unpaired two-tailed *t*-test.

**Fig. S6.**
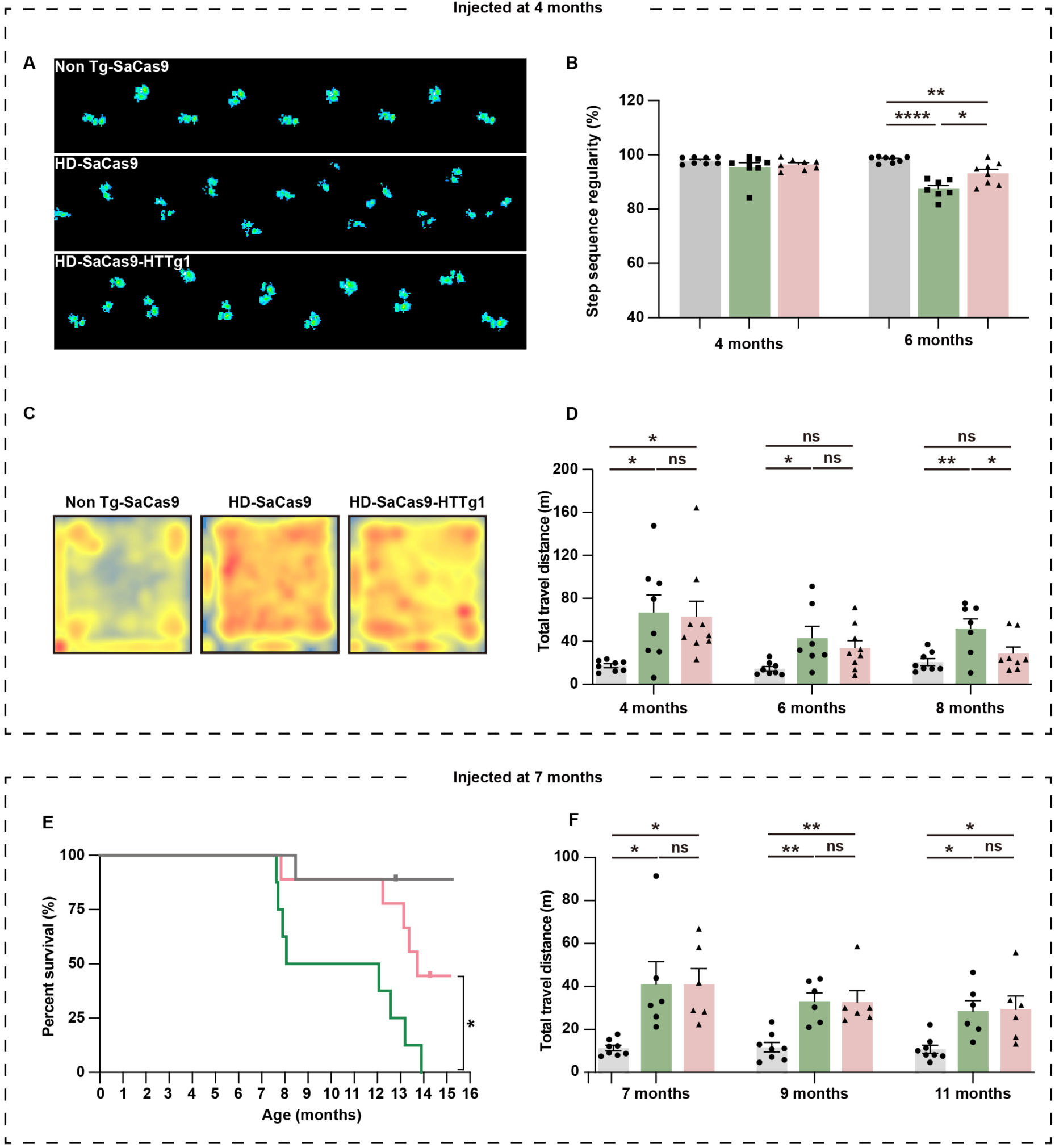
Gene editing at 4 and 7 months resulted in graduated rescue of behavioral phenotypes. (**A** to **D**) Rescue of behavioral performance in 4-month-injected HD-SaCas9-HTTg1 mice. (A and B) Improved gait performance at 6 months. (A) Representative footprints of Non tg-SaCas9 (upper), HD-SaCas9 (middle) and HD-SaCas9-HTTg1 mice (lower). (B) Significant increase of step sequence regularity index in 6-month HD-SaCas9-HTTg1 mice. N=8 mice per group (4 months), N=7 mice (HD-SaCas9, 6 months) and N=8 (Non Tg-SaCas9 and HD-SaCas9-HTTg1, 6 months); \*\*\**p*<0.0001, \*\**p*=0.0066 \**p*=0.0104. (C and D) Partial rescue of locomotor performance in HD-SaCas9-HTTg1 mice in open-field test. (C) Representative heatmaps exhibiting time and location to stay for 8-month-old Non Tg-SaCas9 (left), HD-SaCas9 (middle) and HD-SaCas9-HTTg1 (right) mice. Red color represents cumulative appearance. (D) Total travel distance of 4, 6 and 8-month mice. N=7∼9 mice per group. HD-SaCas9-HTTg1 versus HD-SaCas9, \**p*=0.0488 (8 months); HD-SaCas9-HTTg1 versus Non Tg-SaCas9, \**p*=0.0484 (4 months); HD-SaCas9 versus Non Tg-SaCas9, \*\**p*=0.0073 (8 months), \**p*=0.0339 (6 months), \**p*=0.0359 (4 months). (**E** and **F**) Rescue effects of SaCas9-HTTg1 when injected at 7 months. (E) Partial rescue of survival time in HD-SaCas9-HTTg1 mice compared with HD-SaCas9 (HD-SaCas9-HTTg1 versus HD-SaCas9 *p*=0.0138; HD-SaCas9-HTTg1 versus Non Tg-SaCas9 *p*=0.0959; HD-SaCas9 versus Non Tg-SaCas9 *p*=0.0003). Non Tg-SaCas9 (N=9), HD-SaCas9 (N=8) and HD-SaCas9-HTTg1 (N=9) mice. (F) No significant rescue of locomotor activity in HD-SaCas9-HTTg1 mice in open-field test. N=8 mice (Non Tg-SaCas9), N=6 mice (HD-SaCas9-HTTg1 and HD-SaCas9). HD-SaCas9-HTTg1 versus Non Tg-SaCas9, \**p*=0.0141 (7 months), \*\**p*=0.0023 (9 months), \**p*=0.0162 (11 months); HD-SaCas9 versus Non Tg-SaCas9, \**p*=0.0139 (7 months), \*\**p*=0.0021 (9 months), \**p*=0.0216 (11 months). Data were shown as mean ± SEM. One-way ANOVA with Tukey’s *post hoc* multiple comparisons test (B, D and F) and Log-rank (Mantel-Cox) test (E).

**Fig. S7.**
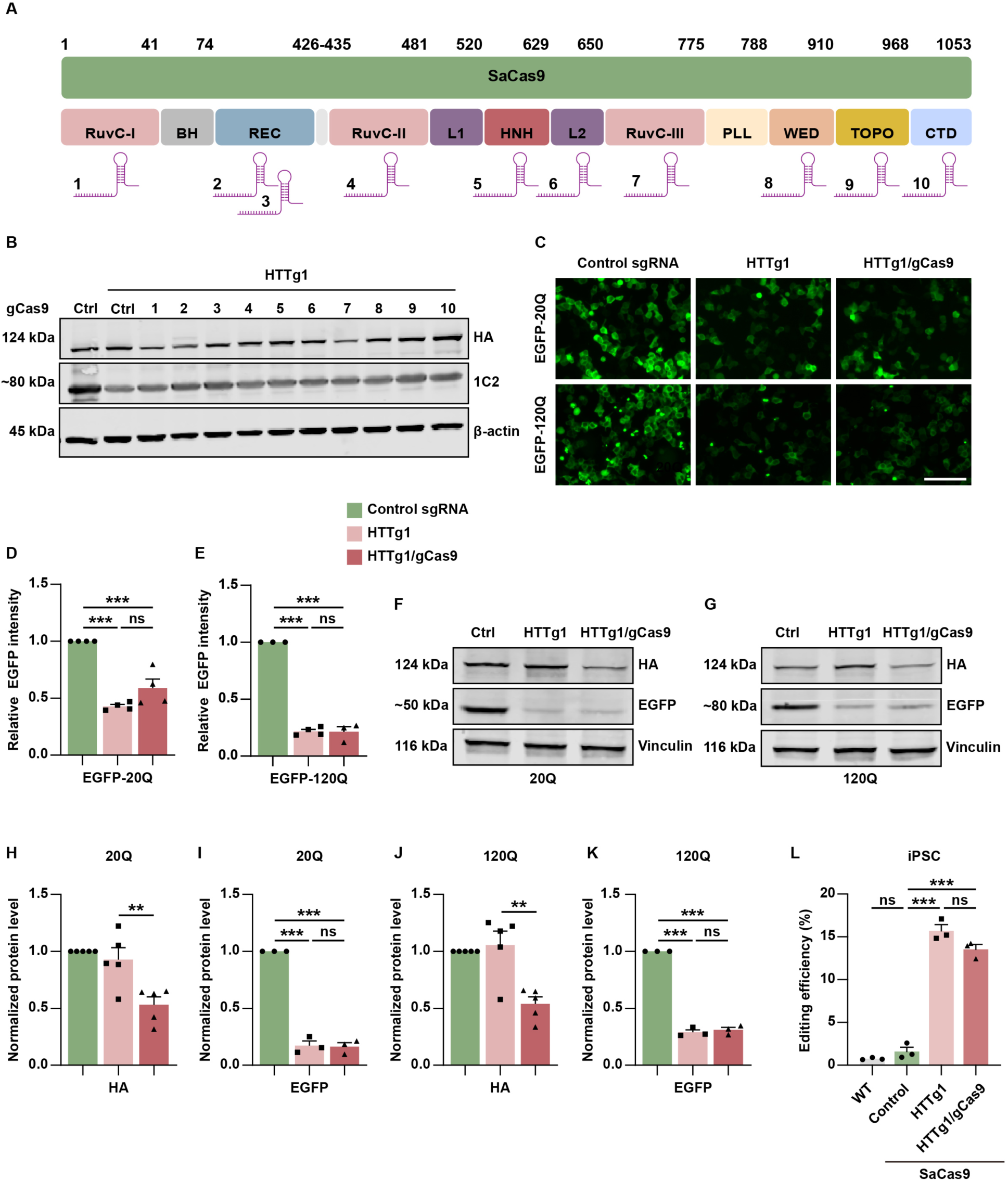
Significant reduction of mHTT and SaCas9 by self-inactivating CRISPR/Cas9 system *in vitro*. (**A**) Schematic position of 10 gCas9 targeting SaCas9. (**B**) In HEK293T cells expressing EGFP-hHTT-120Q, #1 of the 10 tested gCas9 effectively decreased SaCas9 without affecting mHTT reduction. Antibody 1C2 to detect EGFP-hHTT-120Q and anti-HA for SaCas9 detection in Western-blot. (**C** to **E**) HTTg1/gCas9 reduced EGFP-hHTT-20Q/120Q with equal efficiency as HTTg1 in HEK293T cells. (C) Representative images of EGFP fused human HTT exon 1-20Q/120Q. Scale bar: 100 μm. (D) Quantitative analysis of EGFP-20Q fluorescence intensity. N=4 per group; \*\*\**p*<0.0001 (Control sgRNA versus HTTg1), \*\*\**p*=0.0002 (Control sgRNA versus HTTg1/gCas9), ns=0.5469. (E) Quantitative analysis of EGFP-120Q fluorescence intensity. N=4 per group; \*\*\**p*<0.0001 (Control sgRNA versus HTTg1), \*\*\**p*<0.0001 (Control sgRNA versus HTTg1/gCas9), ns=0.9101. (**F** to **K**) HTTg1/gCas9 reduced EGFP-hHTT-20Q/120Q as HTTg1, and decreased SaCas9 in HEK293T cells (F and G). (H and I) Quantification of SaCas9 (anti-HA) expression (H) and EGFP (I) in 20Q. For HA: N=5 per group; \*\**p*=0.0062. For EGFP: N=3 per group; \*\*\**p*<0.0001 (Control sgRNA versus HTTg1), \*\*\**p*<0.0001 (Control sgRNA versus HTTg1/gCas9), ns=0.9818. (J and K) Quantification of SaCas9 (anti-HA) expression (J) and EGFP (K) in 120Q. For HA: N=5 per group; \*\**p*=0.0018. For EGFP: N=3 per group; \*\*\**p*<0.0001 (Control sgRNA versus HTTg1), \*\*\**p*<0.0001 (Control sgRNA versus HTTg1/gCas9), ns=0.7146. (**L**) Editing efficiencies in human iPSC detected by PEM-seq. \*\*\**p*<0.0001, N=3 for each group. Data were shown as mean ± SEM. One-way ANOVA with Tukey’s *post hoc* multiple comparisons test.

**Fig. S8.**
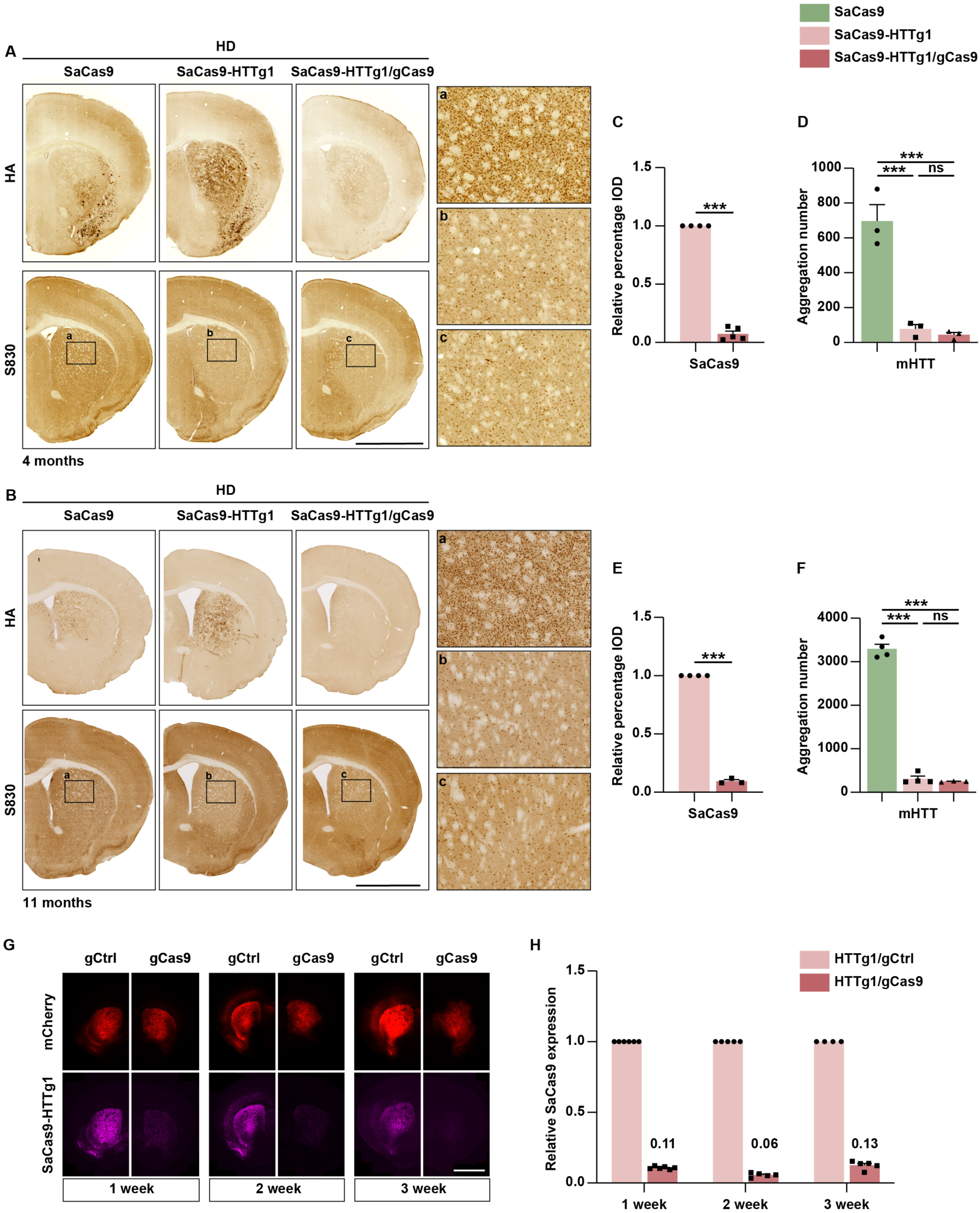
Transient SaCas9 is sufficient for permanent mHTT reduction in self-inactivating CRISPR/Cas9 system. (**A** to **F**) Self-inactivating system decreased the S830-positive mHTT aggregates along with a dramatic reduction of HA-labeled SaCas9. Boxed areas from 4 months old mice (A) and boxed areas from 11 months old mice (B) were enlarged on the right respectively. Scale bar: 2000 μm. (C) Quantification of SaCas9 reduction in a. N=4/5 mice per group; \*\*\**p*<0.0001. (D) Quantification of striatal mHTT aggregation number in boxed areas of (A). N=3 mice per group; \*\*\**p*=0.0006 (HD-SaCas9 versus HD-SaCas9-HTTg1), \*\*\**p*=0.0005 (HD-SaCas9 versus HD-SaCas9-HTTg1/gCas9), ns=0.9084. (E) Quantification of SaCas9 reduction in b. N=4/3 mice per group; \*\*\**p*<0.0001. (F) Quantification of striatal mHTT aggregation number in boxed areas of b. N=4/3 mice per group; \*\*\**p*<0.0001 (HD-SaCas9 versus HD-SaCas9-HTTg1), \*\*\**p*<0.0001 (HD-SaCas9 versus HD-SaCas9-HTTg1/gCas9), ns=0.8507. (**G** and **H**) SaCas9 diminished 1 week post injection with AAV-SaCas9-HTTg1/mCherry-gCas9 (right) compared with AAV-SaCas9-HTTg1/mCherry-gCtrl (left) in mice brain (G). Red signal represented the reporter protein mCherry fluorescence, the purple signal was the expression of SaCas9. Scale bar: 2000 μm. (H) SaCas9 was reduced to 0.11 at week 1, 0.06 at week 2 and 0.13 at week 3 relatively. Data were shown as mean ± SEM. One-way ANOVA with Tukey’s *post hoc* multiple comparisons test and unpaired two-tailed *t*-test.

**Fig. S9.**
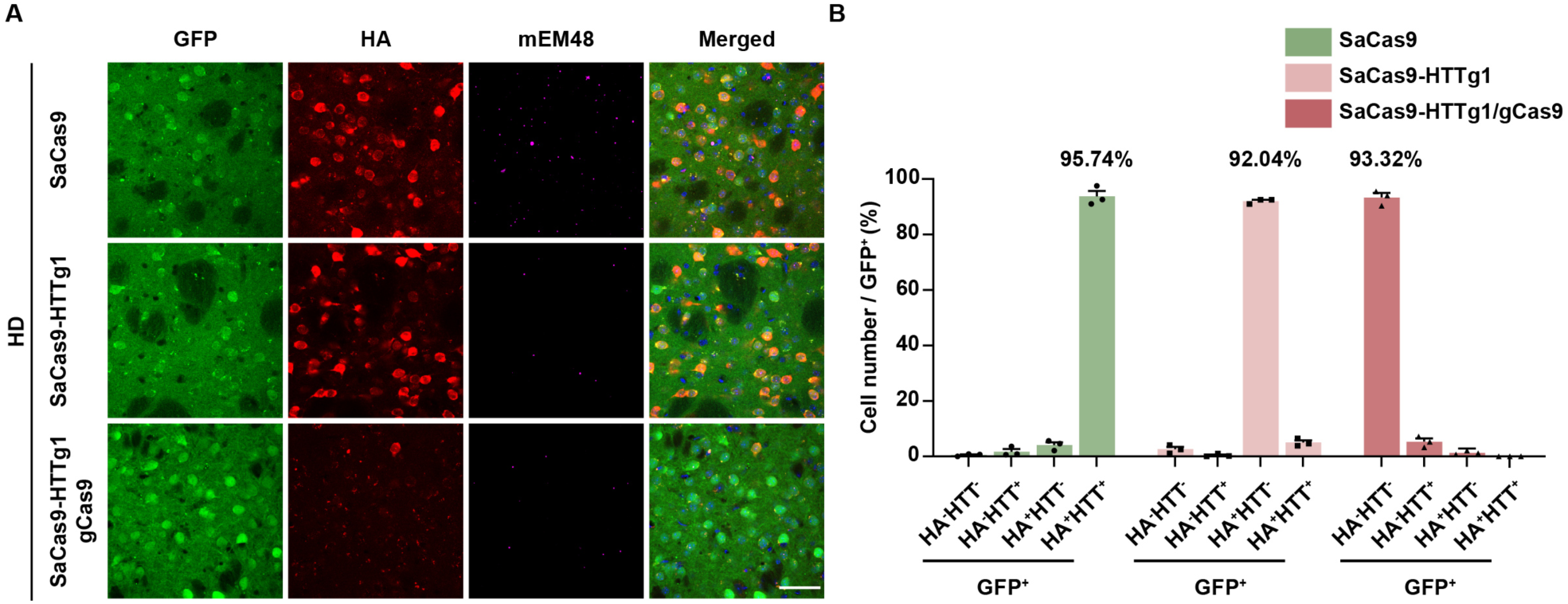
The efficiency of the self-inactivating system on SaCas9 and mHTT reduction. (**A**) Fluorescence schematic of GFP, HA-tagged SaCas9, and mEM48-labeled mHTT aggregates. Scale bar: 50 μm. (**B**) Quantification of HA and mHTT expression in GFP-positive cells from the groups shown in a. In the SaCas9-HTTg1 group, 92% of GFP-positive cells expressed HA but did not express mHTT. In the SaCas9-HTTg1/gCas9 group, 93% of GFP-positive cells expressed neither HA nor mHTT. N = 3 mice per group. data were shown as mean ± SEM.

**Fig. S10.**
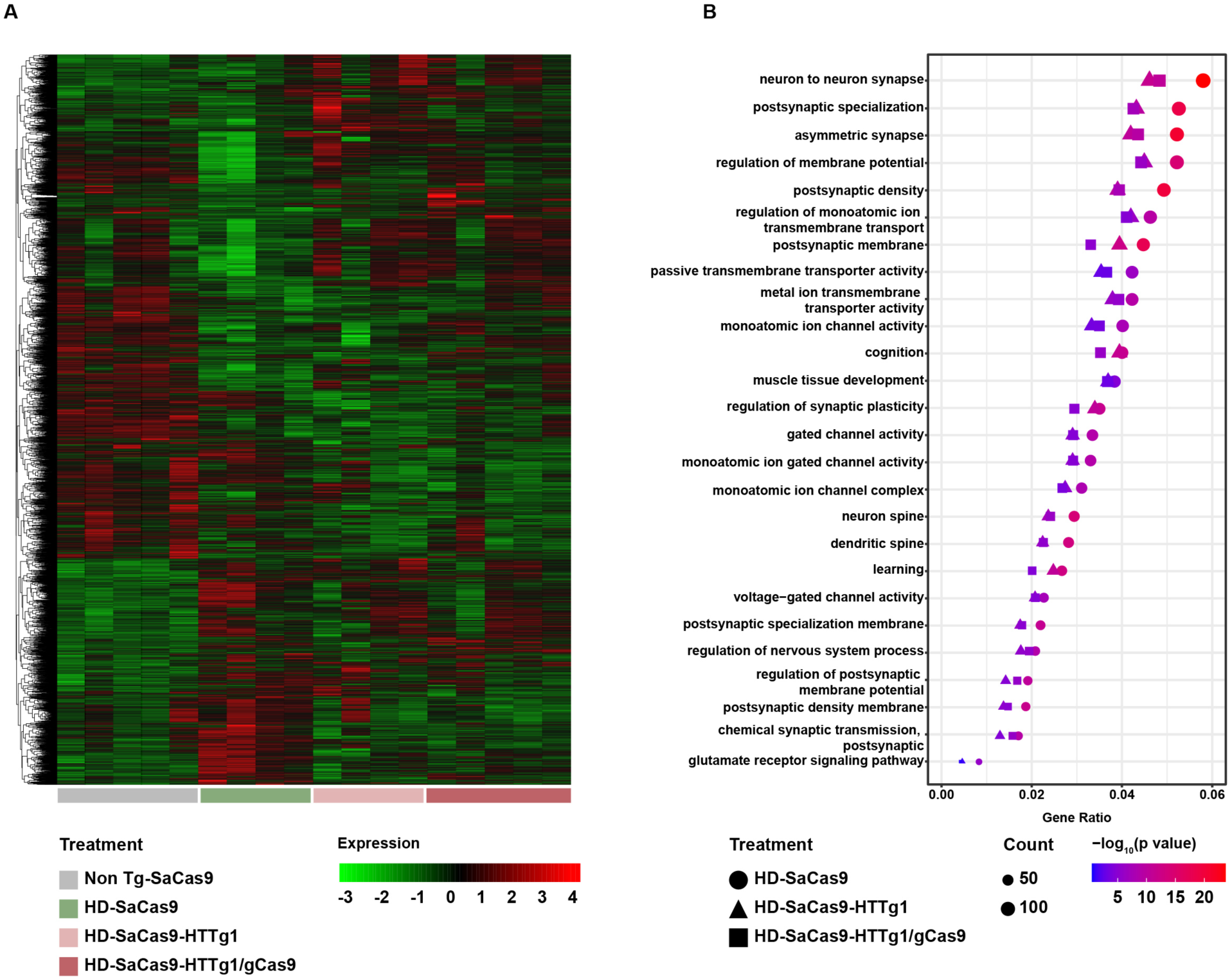
Transcriptomic changes in the striatum of BAC226Q mice following CRISPR/Cas9 gene editing at 1 month of age. (**A**) Heat map of 6,505 gene variance among 4-month-old Non Tg-SaCas9, BAC226Q-SaCas9, BAC226Q-HTTg1 and BAC226Q-HTTg1/gCas9 mice. Green and red color intensities represent gene downregulation and upregulation, respectively. (**B**) GO enrichment of top 26 pathways and the expression of genes enriched in these pathways. RNA-seq analysis revealed that both conventional CRISPR/Cas9 and self-inactivating system ameliorated gene expression dysregulation in BAC226Q mice.

**Table S1.**
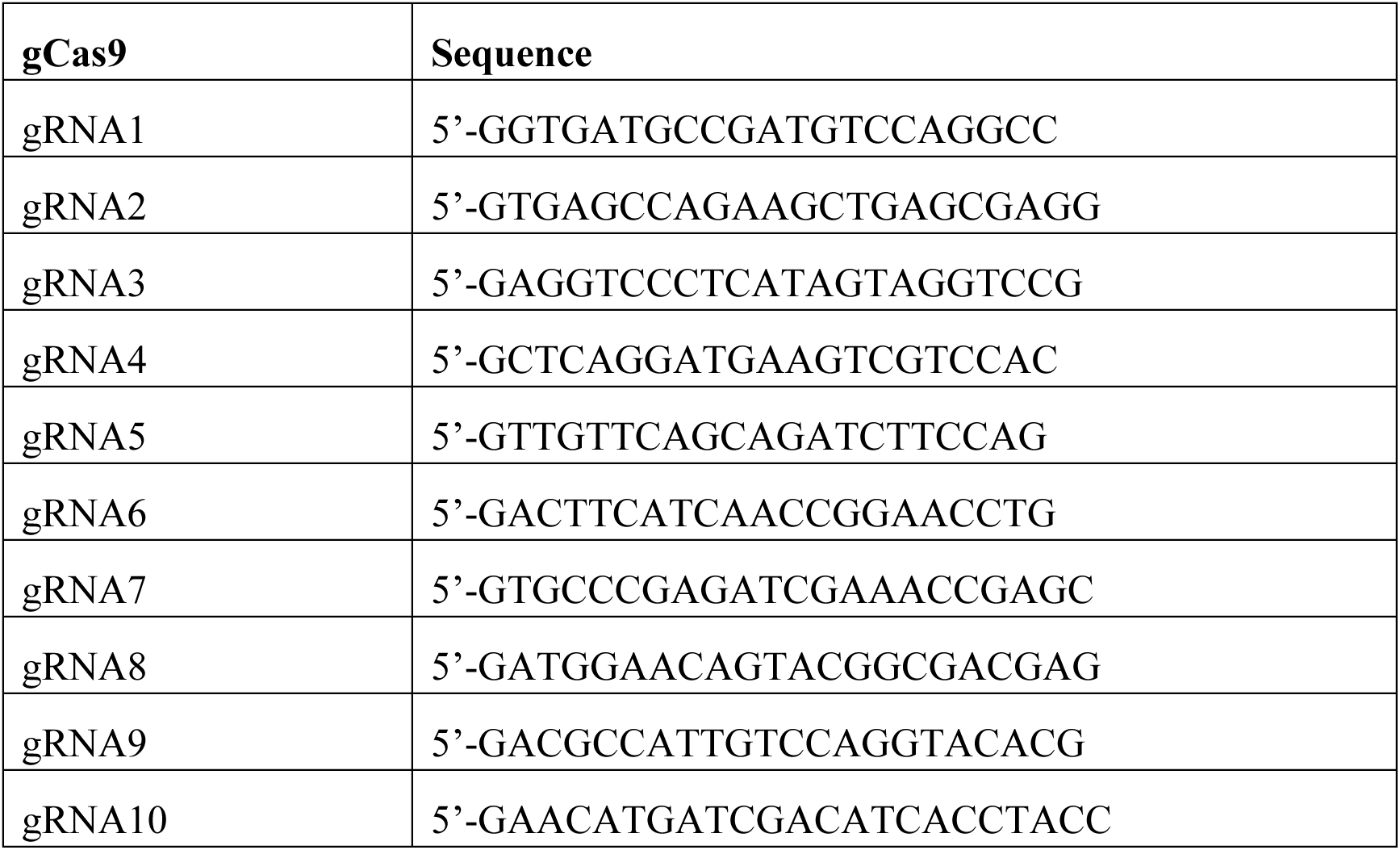
Sequences of 10 gCas9 targeting SaCas9.

### Legends for movies

**Movie S1.** Home cage behavior at 10 months showing abrupt, unnatural jerking and twisting of the head and body in the HD-SaCas9 mouse, improved motor performance in the HD-SaCas9-HTTg1 mouse, and normal behavior in the NonTg-SaCas9 mouse.

**Movie S2.** Improved gait performance in the 6-month-old HD-SaCas9-HTTg1 mouse following gene editing at 1 month of age.

**Movie S3.** Gait improvement in the HD-SaCas9-HTTg1 mouse at 6 months following gene editing at 4 months of age.

**Movie S4.** Improvement of gait performance in a 6-month-old BAC226Q mouse following treatment at 1 month with the self-inactivating AAV-SaCas9-HTTg1/gCas9 system, comparable to conventional CRISPR/Cas9-mediated editing.

## References and Notes

1. GBD 2021 Nervous System Disorders Collaborators, Global, regional, and national burden of disorders affecting the nervous system, 1990-2021: a systematic analysis for the global burden of disease study 2021. Lancet Neurol. 23, 344–381 (2024).

2. L. Gan, M. R. Cookson, L. Petrucelli, A. R. La Spada, Converging pathways in neurodegeneration, from genetics to mechanisms. Nat. Neurosci. 21, 1300–1309 (2018).

3. M. A. Nance, W. Seltzer, T. Ashizawa, R. Bennett, N. McIntosh, R. H. Myers, N. T. Potter, D. K. Shea, Laboratory guidelines for Huntington disease genetic testing. Am. J. Hum. Genet. 62, 1243–1247 (1998).

4. The Huntington’s Disease Collaborative Research Group, A novel gene containing a trinucleotide repeat that is expanded and unstable on Huntington’s disease chromosomes. Cell 72, 971–983 (1993).

5. C. A. Ross, S. J. Tabrizi, Huntington’s disease: from molecular pathogenesis to clinical treatment. Lancet Neurol. 10, 83–98 (2011).

6. F. Squitieri, C. Gellera, M. Cannella, C. Mariotti, G. Cislaghi, D. C. Rubinsztein, E. W. Almqvist, D. Turner, A. C. Bachoud-Lévi, S. A. Simpson, Homozygosity for CAG mutation in Huntington disease is associated with a more severe clinical course. Brain 126, 946–955 (2003).

7. R. K. Graham, E. J. Slow, Y. Deng, N. Bissada, G. Lu, J. Pearson, J. Shehadeh, B. R. Leavitt, L. A. Raymond, M. R. Hayden, Levels of mutant huntingtin influence the phenotypic severity of Huntington disease in YAC128 mouse models. Neurobio. Dis. 21, 444–455 (2006).

8. R. J. Harding, Y. F. Tong, Proteostasis in Huntington’s disease: disease mechanisms and therapeutic opportunities. Acta Pharmacol. Sin. 39, 754–769 (2018).

9. H. B. Kordasiewicz, L. M. Stanek, E. V. Wancewicz, C. Mazur, M. M. McAlonis, K. A. Pytel, J. W. Artates, A. Weiss, S. H. Cheng, L. S. Shihabuddin, Sustained therapeutic reversal of Huntington’s disease by transient repression of huntingtin synthesis. Neuron 74, 1031–1044 (2012).

10. J. B. Carroll, S. C. Warby, A. L. Southwell, C. N. Doty, S. Greenlee, N. Skotte, G. Hung, C. F. Bennett, S. M. Freier, M. R. Hayden, Potent and selective antisense oligonucleotides targeting single-nucleotide polymorphisms in the Huntington disease gene/allele-specific silencing of mutant huntingtin. Mol. Ther. 19, 2178–2185 (2011).

11. A. L. Southwell, H. B. Kordasiewicz, D. Langbehn, N. H. Skotte, M. P. Parsons, E. B. Villanueva, N. S. Caron, M. E. Ostergaard, L. M. Anderson, Y. Xie, L. D. Cengio, H. Findlay-Black, C. N. Doty, B. Fitsimmons, E. E. Swayze, P. P. Seth, L. A. Raymond, C. Frank Bennett, M. R. Hayden, Huntingtin suppression restores cognitive function in a mouse model of Huntington’s disease. Sci. Transl. Med. 10, eaar3959 (2018).

12. S. Q. Harper, P. D. Staber, X. He, S. L. Eliason, I. H. Martins, Q. Mao, L. Yang, R. M. Kotin, H. L. Paulson, B. L. Davidson, RNA interference improves motor and neuropathological abnormalities in a Huntington’s disease mouse model. Proc. Natl. Acad. Sci. USA 102, 5820–5825 (2005).

13. L. M. Stanek, S. P. Sardi, B. Mastis, A. R. Richards, C. M. Treleaven, T. Taksir, K. Misra, S. H. Cheng, L. S. Shihabuddin, Silencing mutant huntingtin by Adeno-associated virus-mediated RNA interference ameliorates disease manifestations in the YAC128 mouse model of Huntington’s disease. Hum. Gene Ther. 25, 461–474 (2014).

14. J. Miniarikova, V. Zimmer, R. Martier, C. C. Brouwers, C. Pythoud, K. Richetin, M. Rey, J. Lubelski, M. M. Evers, S. J. van Deventer, H. Petry, N. Deglon, P. Konstantinova, AAV5-miHTT gene therapy demonstrates suppression of mutant huntingtin aggregation and neuronal dysfunction in a rat model of Huntington’s disease. Gene Ther. 24, 630–639 (2017).

15. A. Valles, M. M. Evers, A. Stam, M. Sogorb-Gonzalez, C. Brouwers, C. Vendrell-Tornero, S. Acar-Broekmans, L. Paerels, J. Klima, B. Bohuslavova, R. Pintauro, V. Fodale, A. Bresciani, R. Liscak, D. Urgosik, Z. Starek, M. Crha, B. Blits, H. Petry, Z. Ellederova, J. Motlik, S. van Deventer, P. Konstantinova, Widespread and sustained target engagement in Huntington’s disease minipigs upon intrastriatal microRNA-based gene therapy. Sci. Transl. Med. 13, eabb8920 (2021).

16. S. B. Thomson, A. Stam, C. Brouwers, V. Fodale, A. Bresciani, M. Vermeulen, S. Mostafavi, T. L. Petkau, A. Hill, A. Yung, B. Russell-Schulz, P. Kozlowski, A. MacKay, D. Ma, M. F. Beg, M. M. Evers, A. Valles, B. R. Leavitt, AAV5-miHTT-mediated huntingtin lowering improves brain health in a Huntington’s disease mouse model. Brain 146, 2298–2315 (2023).

17. L. Zhang, T. Wu, Y. Shan, G. Li, X. Ni, X. Chen, X. Hu, L. Lin, Y. Li, Y. Guan, J. Gao, D. Chen, Y. Zhang, Z. Pei, X. Chen, Therapeutic reversal of Huntington’s disease by in vivo self-assembled siRNAs. Brain 144, 3421–3435 (2021).

18. J. L. McBride, M. R. Pitzer, R. L. Boudreau, B. Dufour, T. Hobbs, S. R. Ojeda, B. L. Davidson, Preclinical safety of RNAi-mediated HTT suppression in the rhesus macaque as a potential therapy for Huntington’s disease. Mol. Ther. 19, 2152–2162 (2011).

19. J. E. Powell, C. K. W. Lim, R. Krishnan, T. X. McCallister, C. Saporito-Magrina, M. A. Zeballos, G. D. McPheron, T. Gaj, Targeted gene silencing in the nervous system with CRISPR-Cas13. Sci. Adv. 8, eabk2485 (2022).

20. K. H. Morelli, Q. Wu, M. L. Gosztyla, H. Liu, M. Yao, C. Zhang, J. Chen, R. J. Marina, K. Lee, K. L. Jones, M. Y. Huang, A. Li, C. Smith-Geater, L. M. Thompson, W. Duan, G. W. Yeo, An RNA-targeting CRISPR-Cas13d system alleviates disease-related phenotypes in Huntington’s disease models. Nat. Neurosci. 26, 27–38 (2023).

21. S. J. Tabrizi, B. R. Leavitt, G. B. Landwehrmeyer, E. J. Wild, C. Saft, R. A. Barker, N. F. Blair, D. Craufurd, J. Priller, H. Rickards, A. Rosser, H. B. Kordasiewicz, C. Czech, E. E. Swayze, D. A. Norris, T. Baumann, I. Gerlach, S. A. Schobel, E. Paz, A. V. Smith, C. F. Bennett, R. M. Lane, I.-H. S. S. T. Phase 1-2a, Targeting huntingtin expression in patients with Huntington’s disease. N. Engl. J. Med. 380, 2307–2316 (2019).

22. M. Garriga-Canut, C. Agustin-Pavon, F. Herrmann, A. Sanchez, M. Dierssen, C. Fillat, M. Isalan, Synthetic zinc finger repressors reduce mutant huntingtin expression in the brain of R6/2 mice. Proc. Natl. Acad. Sci. USA 109, 3136–3145 (2012).

23. B. Zeitler, S. Froelich, K. Marlen, D. A. Shivak, Q. Yu, D. Li, J. R. Pearl, J. C. Miller, L. Zhang, D. E. Paschon, S. J. Hinkley, I. Ankoudinova, S. Lam, D. Guschin, L. Kopan, J. M. Cherone, H. B. Nguyen, G. Qiao, Y. Ataei, M. C. Mendel, R. Amora, R. Surosky, J. Laganiere, B. J. Vu, A. Narayanan, Y. Sedaghat, K. Tillack, C. Thiede, A. Gartner, S. Kwak, J. Bard, L. Mrzljak, L. Park, T. Heikkinen, K. K. Lehtimaki, M. M. Svedberg, J. Haggkvist, L. Tari, M. Toth, A. Varrone, C. Halldin, A. E. Kudwa, S. Ramboz, M. Day, J. Kondapalli, D. J. Surmeier, F. D. Urnov, P. D. Gregory, E. J. Rebar, I. Munoz-Sanjuan, H. S. Zhang, Allele-selective transcriptional repression of mutant HTT for the treatment of Huntington’s disease. Nat. Med. 25, 1131–1142 (2019).

24. C. Agustin-Pavon, M. Mielcarek, M. Garriga-Canut, M. Isalan, Deimmunization for gene therapy: host matching of synthetic zinc finger constructs enables long-term mutant Huntingtin repression in mice. Mol. Neurodegener. 11, 64 (2016).

25. J. W. Shin, K. H. Kim, M. J. Chao, R. S. Atwal, T. Gillis, M. E. MacDonald, J. F. Gusella, J. M. Lee, Permanent inactivation of Huntington’s disease mutation by personalized allele-specific CRISPR/Cas9. Hum. Mol. Genet. 25, 4566–4576 (2016).

26. A. M. Monteys, S. A. Ebanks, M. S. Keiser, B. L. Davidson, CRISPR/Cas9 editing of the mutant huntingtin allele in vitro and in vivo. Mol. Ther. 25, 12–23 (2017).

27. S. R. Oikemus, E. L. Pfister, E. Sapp, K. O. Chase, L. A. Kennington, E. Hudgens, R. Miller, L. J. Zhu, A. Chaudhary, E. O. Mick, M. Sena-Esteves, S. A. Wolfe, M. DiFiglia, N. Aronin, M. H. Brodsky, Allele-specific knockdown of mutant huntingtin protein via editing at coding region single nucleotide polymorphism heterozygosities. Hum. Gene Ther. 33, 25–36 (2022).

28. M. Dabrowska, W. Juzwa, W. J. Krzyzosiak, M. Olejniczak, Precise excision of the CAG tract from the huntingtin gene by Cas9 nickases. Front. Neurosci. 12, 75 (2018).

29. M. Jinek, K. Chylinski, I. Fonfara, M. Hauer, J. A. Doudna, E. Charpentier, A programmable dual-RNA-guided DNA endonuclease in adaptive bacterial immunity. Science 337, 816–821 (2012).

30. L. Cong, F. A. Ran, D. Cox, S. Lin, R. Barretto, N. Habib, P. D. Hsu, X. Wu, W. Jiang, L. A. Marraffini, F. Zhang, Multiplex genome engineering using CRISPR/Cas systems. Science 339, 819–823 (2013).

31. P. Mali, L. Yang, K. M. Esvelt, J. Aach, M. Guell, J. E. DiCarlo, J. E. Norville, G. M. Church, RNA-guided human genome engineering via Cas9. Science 339, 823–826 (2013).

32. M. Jinek, A. East, A. Cheng, S. Lin, E. Ma, J. Doudna, RNA-programmed genome editing in human cells. Elife 2, e00471 (2013).

33. A. Li, C. M. Lee, A. E. Hurley, K. E. Jarrett, M. De Giorgi, W. Lu, K. S. Balderrama, A. M. Doerfler, H. Deshmukh, A. Ray, G. Bao, W. R. Lagor, A self-deleting AAV-CRISPR system for in vivo genome editing. Mol. Ther. Methods Clin. Dev. 12, 111–122 (2019).

34. N. Merienne, G. Vachey, L. de Longprez, C. Meunier, V. Zimmer, G. Perriard, M. Canales, A. Mathias, L. Herrgott, T. Beltraminelli, A. Maulet, T. Dequesne, C. Pythoud, M. Rey, L. Pellerin, E. Brouillet, A. L. Perrier, R. du Pasquier, N. Deglon, The self-inactivating KamiCas9 system for the editing of CNS disease genes. Cell Rep. 20, 2980–2991 (2017).

35. Y. Chen, X. Liu, Y. Zhang, H. Wang, H. Ying, M. Liu, D. Li, K. O. Lui, Q. Ding, A self-restricted CRISPR system to reduce off-target effects. Mol. Ther. 24, 1508–1510 (2016).

36. G. Petris, A. Casini, C. Montagna, F. Lorenzin, D. Prandi, A. Romanel, J. Zasso, L. Conti, F. Demichelis, A. Cereseto, Hit and go Cas9 delivered through a lentiviral based self-limiting circuit. Nat. Commun. 8, 15334 (2017).

37. A. M. Monteys, A. A. Hundley, P. T. Ranum, L. Tecedor, A. Muehlmatt, E. Lim, D. Lukashev, R. Sivasankaran, B. L. Davidson, Regulated control of gene therapies by drug-induced splicing. Nature 596, 291–295 (2021).

38. S. Yang, R. Chang, H. Yang, T. Zhao, Y. Hong, H. E. Kong, X. Sun, Z. Qin, P. Jin, S. Li, X. J. Li, CRISPR/Cas9-mediated gene editing ameliorates neurotoxicity in mouse model of Huntington’s disease. J. Clin. Invest. 127, 2719–2724 (2017).

39. F. K. Ekman, D. S. Ojala, M. M. Adil, P. A. Lopez, D. V. Schaffer, T. Gaj, CRISPR-Cas9-mediated genome editing increases lifespan and improves motor deficits in a Huntington’s disease mouse model. Mol. Ther. Nucleic Acids 17, 829–839 (2019).

40. S. Yan, X. Zheng, Y. Q. Lin, C. J. Li, Z. M. Liu, J. W. Li, Z. C. Tu, Y. Zhao, C. H. Huang, Y. Z. Chen, J. Li, X. C. Song, B. F. Han, W. Wang, W. E. Liang, L. X. Lai, X. J. Li, S. H. Li, Cas9-mediated replacement of expanded CAG repeats in a pig model of Huntington’s disease. Nat. Biomed. Eng. 7, 629-+ (2023).

41. E. A. Spronck, C. C. Brouwers, A. Valles, M. de Haan, H. Petry, S. J. van Deventer, P. Konstantinova, M. M. Evers, AAV5-miHTT gene therapy demonstrates sustained huntingtin lowering and functional improvement in Huntington disease mouse models. Mol. Ther. Methods Clin. Dev. 13, 334–343 (2019).

42. G. Wang, X. Liu, M. A. Gaertig, S. Li, X. J. Li, Ablation of huntingtin in adult neurons is nondeleterious but its depletion in young mice causes acute pancreatitis. Proc. Natl. Acad. Sci. USA 113, 3359–3364 (2016).

43. R. Grondin, M. D. Kaytor, Y. Ai, P. T. Nelson, D. R. Thakker, J. Heisel, M. R. Weatherspoon, J. L. Blum, E. N. Burright, Z. Zhang, W. F. Kaemmerer, Six-month partial suppression of Huntingtin is well tolerated in the adult rhesus striatum. Brain 135, 1197–1209 (2012).

44. P. Dietrich, I. M. Johnson, S. Alli, I. Dragatsis, Elimination of huntingtin in the adult mouse leads to progressive behavioral deficits, bilateral thalamic calcification, and altered brain iron homeostasis. PLoS Genet. 13, e1006846 (2017).

45. S. Regio, G. Vachey, E. Goni, F. Duarte, M. Rybarikova, M. Sipion, M. Rey, M. Huarte, N. Deglon, Revisiting the outcome of adult wild-type Htt inactivation in the context of HTT-lowering strategies for Huntington’s disease. Brain Commun. 5, fcad344 (2023).

46. R. E. Handsaker, S. Kashin, N. M. Reed, S. Tan, W. S. Lee, T. M. McDonald, K. Morris, N. Kamitaki, C. D. Mullally, N. R. Morakabati, M. Goldman, G. Lind, R. Kohli, E. Lawton, M. Hogan, K. Ichihara, S. Berretta, S. A. McCarroll, Long somatic DNA-repeat expansion drives neurodegeneration in Huntington’s disease. Cell 188, 623–639 e619 (2025).

47. N. Wang, S. Zhang, P. Langfelder, L. Ramanathan, F. Gao, M. Plascencia, R. Vaca, X. Gu, L. Deng, L. E. Dionisio, H. Vu, E. Maciejewski, J. Ernst, B. C. Prasad, T. F. Vogt, S. Horvath, J. S. Aaronson, J. Rosinski, X. W. Yang, Distinct mismatch-repair complex genes set neuronal CAG-repeat expansion rate to drive selective pathogenesis in HD mice. Cell 188, 1524–1544 e1522 (2025).

48. S. A. Shenoy, S. Zheng, W. Liu, Y. Dai, Y. Liu, Z. Hou, S. Mori, Y. Tang, J. Cheng, W. Duan, C. Li, A novel and accurate full-length HTT mouse model for Huntington’s disease. Elife 11, e70217 (2022).

49. X. Gu, J. P. Cantle, E. R. Greiner, C. Y. Lee, A. M. Barth, F. Gao, C. S. Park, Z. Zhang, S. Sandoval-Miller, R. L. Zhang, M. Diamond, I. Mody, G. Coppola, X. W. Yang, N17 modifies mutant Huntingtin nuclear pathogenesis and severity of disease in HD BAC transgenic mice. Neuron 85, 726–741 (2015).

50. M. DiFiglia, E. Sapp, K. O. Chase, S. W. Davies, G. P. Bates, J. P. Vonsattel, N. Aronin, Aggregation of huntingtin in neuronal intranuclear inclusions and dystrophic neurites in brain. Science 277, 1990–1993 (1997).

51. D. M. Marchionini, J.-P. Liu, A. Ambesi-Impiombato, K. Kerker, K. Cirillo, M. Bansal, R. Mushlin, D. Brunner, S. Ramboz, M. Kwan, K. Kuhlbrodt, K. Tillack, F. Peters, L. Rauhala, J. Obenauer, J. R. Greene, C. Hartl, V. Khetarpal, B. Lager, J. Rosinski, J. Aaronson, M. Alam, E. Signer, I. Muñoz-Sanjuán, D. Howland, S. O. Zeitlin, Benefits of global mutant huntingtin lowering diminish over time in a Huntington’s disease mouse model. JCI Insight 7, (2022).

52. P. Langfelder, J. P. Cantle, D. Chatzopoulou, N. Wang, F. Gao, I. Al-Ramahi, X. H. Lu, E. M. Ramos, K. El-Zein, Y. Zhao, S. Deverasetty, A. Tebbe, C. Schaab, D. J. Lavery, D. Howland, S. Kwak, J. Botas, J. S. Aaronson, J. Rosinski, G. Coppola, S. Horvath, X. W. Yang, Integrated genomics and proteomics define huntingtin CAG length-dependent networks in mice. Nat. Neurosci. 19, 623–633 (2016).

53. F. A. Ran, L. Cong, W. X. Yan, D. A. Scott, J. S. Gootenberg, A. J. Kriz, B. Zetsche, O. Shalem, X. Wu, K. S. Makarova, E. V. Koonin, P. A. Sharp, F. Zhang, In vivo genome editing using Staphylococcus aureus Cas9. Nature 520, 186–191 (2015).

54. J. Yin, M. Liu, Y. Liu, J. Wu, T. Gan, W. Zhang, Y. Li, Y. Zhou, J. Hu, Optimizing genome editing strategy by primer-extension-mediated sequencing. Cell Discov. 5, 18 (2019).

55. G. P. Bates, R. Dorsey, J. F. Gusella, M. R. Hayden, C. Kay, B. R. Leavitt, M. Nance, C. A. Ross, R. I. Scahill, R. Wetzel, E. J. Wild, S. J. Tabrizi, Huntington disease. Nat. Rev. Dis. Primers 1, 15005 (2015).

56. N. Wang, M. Gray, X. H. Lu, J. P. Cantle, S. M. Holley, E. Greiner, X. Gu, D. Shirasaki, C. Cepeda, Y. Li, H. Dong, M. S. Levine, X. W. Yang, Neuronal targets for reducing mutant huntingtin expression to ameliorate disease in a mouse model of Huntington’s disease. Nat. Med. 20, 536–541 (2014).

57. E. Hudry, L. H. Vandenberghe, Therapeutic AAV gene transfer to the nervous system: a clinical reality. Neuron 101, 839–862 (2019).

58. S. N. Mathiesen, J. L. Lock, L. Schoderboeck, W. C. Abraham, S. M. Hughes, CNS transduction benefits of AAV-PHP.eB over AAV9 are dependent on administration route and mouse strain. Mol. Ther. Methods Clin. Dev. 19, 447–458 (2020).

59. C. Hinderer, N. Katz, E. L. Buza, C. Dyer, T. Goode, P. Bell, L. K. Richman, J. M. Wilson, Severe toxicity in nonhuman primates and piglets following high-dose intravenous administration of an Adeno-associated virus vector expressing human SMN. Hum. Gene Ther. 29, 285–298 (2018).

60. A. Lek, B. Wong, A. Keeler, M. Blackwood, K. Ma, S. Huang, K. Sylvia, A. R. Batista, R. Artinian, D. Kokoski, S. Parajuli, J. Putra, C. K. Carreon, H. Lidov, K. Woodman, S. Pajusalu, J. M. Spinazzola, T. Gallagher, J. LaRovere, D. Balderson, L. Black, K. Sutton, R. Horgan, M. Lek, T. Flotte, Death after high-dose rAAV9 gene therapy in a patient with Duchenne’s Muscular Dystrophy. N. Eng. J. Med. 389, 1203–1210 (2023).

61. Y. K. Chan, S. K. Wang, C. J. Chu, D. A. Copland, A. J. Letizia, H. Costa Verdera, J. J. Chiang, M. Sethi, M. K. Wang, W. J. Neidermyer, Y. Chan, E. T. Lim, A. R. Graveline, M. Sanchez, R. F. Boyd, T. S. Vihtelic, R. G. C. O. Inciong, J. M. Slain, P. J. Alphonse, Y. Xue, L. R. Robinson-McCarthy, J. M. Tam, M. H. Jabbar, B. Sahu, J. F. Adeniran, M. Muhuri, P. W. L. Tai, J. Xie, T. B. Krause, A. Vernet, M. Pezone, R. Xiao, T. Liu, W. Wang, H. J. Kaplan, G. Gao, A. D. Dick, F. Mingozzi, M. A. McCall, C. L. Cepko, G. M. Church, Engineering adeno-associated viral vectors to evade innate immune and inflammatory responses. Sci. Transl. Med. 13, eabd3438 (2021).

62. J. Bradford, J. Y. Shin, M. Roberts, C. E. Wang, X. J. Li, S. Li, Expression of mutant huntingtin in mouse brain astrocytes causes age-dependent neurological symptoms. Proc. Natl. Acad. Sci. USA 106, 22480–22485 (2009).

63. A. Benraiss, S. Wang, S. Herrlinger, X. J. Li, D. Chandler-Militello, J. Mauceri, H. B. Burm, M. Toner, M. Osipovitch, Q. W. J. Xu, F. F. Ding, F. S. Wang, N. Kang, J. Kang, P. C. Curtin, D. Brunner, M. S. Windrem, I. Munoz-Sanjuan, M. Nedergaard, S. A. Goldman, Human glia can both induce and rescue aspects of disease phenotype in Huntington disease. Nat. Commun. 7, 11758 (2016).

64. M. Faideau, J. Kim, K. Cormier, R. Gilmore, M. Welch, G. Auregan, N. Dufour, M. Guillermier, E. Brouillet, P. Hantraye, N. Deglon, R. J. Ferrante, G. Bonvento, In vivo expression of polyglutamine-expanded huntingtin by mouse striatal astrocytes impairs glutamate transport: a correlation with Huntington’s disease subjects. Hum. Mol. Genet. 19, 3053–3067 (2010).

65. Y. Hong, T. Zhao, X. J. Li, S. H. Li, Mutant huntingtin impairs BDNF release from astrocytes by disrupting conversion of Rab3a-GTP into Rab3a-GDP. J. Neurosci. 36, 8790–8801 (2016).

66. S. J. Tabrizi, S. Schobel, E. C. Gantman, A. Mansbach, B. Borowsky, P. Konstantinova, T. A. Mestre, J. Panagoulias, C. A. Ross, M. Zauderer, A. P. Mullin, K. Romero, S. Sivakumaran, E. C. Turner, J. D. Long, C. Sampaio, C. Huntington’s Disease Regulatory Science, A biological classification of Huntington’s disease: the Integrated Staging System. Lancet Neurol. 21, 632–644 (2022).

67. D. W. Morgens, M. Wainberg, E. A. Boyle, O. Ursu, C. L. Araya, C. K. Tsui, M. S. Haney, G. T. Hess, K. Han, E. E. Jeng, A. Li, M. P. Snyder, W. J. Greenleaf, A. Kundaje, M. C. Bassik, Genome-scale measurement of off-target activity using Cas9 toxicity in high-throughput screens. Nat. Commun. 8, 15178 (2017).

68. J. P. K. Bravo, M.-S. Liu, G. N. Hibshman, T. L. Dangerfield, K. Jung, R. S. McCool, K. A. Johnson, D. W. Taylor, Structural basis for mismatch surveillance by CRISPR–Cas9. Nature 603, 343–347 (2022).

69. J. Yin, R. Lu, C. Xin, Y. Wang, X. Ling, D. Li, W. Zhang, M. Liu, W. Xie, L. Kong, W. Si, P. Wei, B. Xiao, H.-Y. Lee, T. Liu, J. Hu, Cas9 exo-endonuclease eliminates chromosomal translocations during genome editing. Nat. Commun. 13, 1204 (2022).

70. C. T. Charlesworth, P. S. Deshpande, D. P. Dever, J. Camarena, V. T. Lemgart, M. K. Cromer, C. A. Vakulskas, M. A. Collingwood, L. Zhang, N. M. Bode, M. A. Behlke, B. Dejene, B. Cieniewicz, R. Romano, B. J. Lesch, N. Gomez-Ospina, S. Mantri, M. Pavel-Dinu, K. I. Weinberg, M. H. Porteus, Identification of preexisting adaptive immunity to Cas9 proteins in humans. Nat. Med. 25, 249–254 (2019).

71. W. Ge, S. Gou, X. Zhao, Q. Jin, Z. Zhuang, Y. Zhao, Y. Liang, Z. Ouyang, X. Liu, F. Chen, H. Shi, H. Yan, H. Wu, L. Lai, K. Wang, In vivo evaluation of guide-free Cas9-induced safety risks in a pig model. Signal Transduc. Target. Ther. 9, 184 (2024).

72. E. A. Kappos, P. K. Sieber, P. E. Engels, A. V. Mariolo, S. D’Arpa, D. J. Schaefer, D. F. Kalbermatten, Validity and reliability of the CatWalk system as a static and dynamic gait analysis tool for the assessment of functional nerve recovery in small animal models. Brain Behav. 7, e00723 (2017).

